# Accumulation of two distinct alleles causes astringency loss in cultivated polyploid persimmon

**DOI:** 10.64898/2026.10.31.762479

**Authors:** Yosuke Fujiwara, Soichiro Nishiyama, Noriyuki Onoue, Ryusuke Matsuzaki, Atsushi Kono, Akifumi Azuma, Toshihiro Saito, Takeo Shimizu, Akihiko Sato, Takuji Ichino, Hisayo Yamane, Keizo Yonemori, Ryutaro Tao

## Abstract

Persimmon fruit accumulates high levels of proanthocyanidins (PAs), causing extreme astringency. In pollination-constant and non-astringent (PCNA) cultivars, PA accumulation ceases early in fruit development. Although the PCNA trait is recessively controlled by the *ASTRINGENCY* (*AST*) locus, its causal gene has remained unknown. Using haplotype-resolved forward genetics in a hexaploid genetic background, we identified the previously uncharacterized *TANNIN* (*TAN*) as the causal gene. Long-read sequencing revealed that PCNA originated through the accumulation of independently derived *ast* alleles that cause transcriptional deficiency of *TAN*. These findings resolve the complex genetic basis of natural astringency loss in persimmon and provide a framework for molecular breeding. Further investigation of TAN function would uncover mechanisms underlying massive accumulation of PAs in persimmon fruit and insight into specialized metabolite sequestration.

## Introduction

Proanthocyaindins (PAs), also known as condensed tannins, are widespread specialized metabolites in land plants. PAs strongly influence food quality by causing astringency, while also providing potential health benefits through their antioxidant properties and microbiota modulation^1, 2^. PAs are polymers of flavan-3-ols and are highly reactive and potentially cytotoxic; therefore, their intracellular sequestration is critical for maintaining cellular integrity. Although there have been much progress in understanding the molecular basis of PA biosynthesis, the mechanisms underlying PA polymerization and intracellular trafficking into the vacuole remain unresolved^3, 4^. This gap limits our understanding of how plants achieve the quantitative control of PA accumulation.

Persimmon (*Diospyros kaki* Thunb.; 2*n* = 6*x* = 90) is a fruit tree crop that originated in east Asia. A unique characteristic of persimmon is the massive accumulation of PAs, which causes strong astringency of the fruit. Persimmon cultivars have long been classified into pollination-constant and astringent (PCA), pollination-variant and astringent (PVA), pollination-variant and non-astringent (PVNA) and pollination-constant and non-astringent (PCNA)^5, 6^. In non-PCNA cultivars, astringency loss in mature fruit is associated with seed-derived volatile compounds^7^ and is quantitatively regulated. On the other hand, PCNA cultivars are unique in that PA accumulation ceases early during fruit development, and genes related to PA biosynthesis decrease coinciding with the cessation of PA accumulation^5, 8, 9^. PCNA phenotype is conferred by a single locus called *ASTRINGENCY* (*AST*), and PCNA is recessive homozygous (*aaaaaa*)^10, 11^. This natural loss of astringency is a commercially essential trait in modern persimmon breeding. However, PCNA germplasm remains rare and genetically narrow, consistent with their presumed recent origin from a limited founder background^12^. Elucidation of the causal gene will therefore provide both a foundation for molecular breeding and a unique framework for dissecting mechanisms underlying continuous and massive PA accumulation. An *ast*-linked insertion, Indel-3, has been found and intensively used for marker-assisted selection of PCNA offspring^13, 14^. However, the causal gene of *AST* locus has not been identified due to the complexity of polyploid persimmon genome. We previously delimited the *AST* region using a segregating population of hexaploid *D. kaki* and found that the corresponding interval spans 915 kb, including Indel-3, in the diploid *D. lotus* (2*n* = 2*x* =30) genome^15^. However, no gene within this interval showed clear association with PA biosynthesis or differential expression between PCNA and non-PCNA genotypes^15, 16^. These results suggested that a haplotype-resolved analysis of the hexaploid *D. kaki* genome is required to fully characterize the candidate genes and allelic variations at the *AST* locus.

## Results

### Identification of a single candidate responsible for PCNA trait by haplotype-resolved hexaploid genome analysis

We first established haplotype-resolved sequences covering the delimited *AST* region in the non-PCNA cultivar ‘Taiten’. Previous analyses revealed that ‘Taiten’ carries six distinct haplotypes (*A*224, *A*240, *a*349, *a*353_2a, *a*353_2b, and *a*355) at the Indel-3 region^17^. To enable accurate haplotype reconstruction, we obtained 43.5X coverage of Pacbio HiFi reads from ‘Taiten’ and applied a trio-binning strategy, in which HiFi reads are partitioned into each parental group prior to assembly^18^ (Supplementary Fig. 1, Supplementary Table 1). In our study, based on partitioning accuracy test of each parent-specific k-mer length (Supplementary Table 2), whole-genome reads were partitioned into three groups: hapA (derived from ‘Kurokuma’; *A*224, *A*240 and *a*353_2a), hapB (derived from ‘Taishu’; *a*349, *a*353_2b and *a*355), and hapU (unclassified).

Using k = 25 and 31 partitioning results, de novo assemblies were performed separately for each parental group (Supplementary Table 3, Supplementary Table 4). As a result, assembly in k = 25 failed to generate complete haplotype sequences (Supplementary Fig. 2). In k = 31, five contigs covering the region were produced, and *A*224, *A*240, and *a*353_2a were generated from the ‘Kurokuma’ assembly, whereas *a*349, *a*353_2a, and *a*353_2b were generated from the ‘Taishu’ assembly (Fig. 1A). Notably, polymorphisms used for the *A*224- and *A*240-specific markers^15^ were confirmed in the corresponding contigs and were consistent across the delimited region (Fig. 1A). However, *a*353_2a was assembled from both parental read groups, and *a*355 was not assembled, indicating incomplete partitioning of reads specific to this haplotype under the current parameters. Previous studies showed that *a*355 is highly similar to *a*353_2b, differing only in short sequence repeat (SSR) length^17^. Consistent with this, no sequence polymorphisms were detected between *a*353_2b and *a*355 within the HiFi read span (∼18 kb), except for the SSR (Supplementary Fig. 3). Therefore, we classified *a*353 and *a*355 into the same haplotype group for the following analysis. Nevertheless, the k = 31 assembly resolved all sequence-distinguishable *AST* haplotypes as independent contigs spanning the delimited region. Moreover, no large structural variant was detected in these contigs compared to the diploid *D. oleifera* sequence^19^.

**Figure 1.**
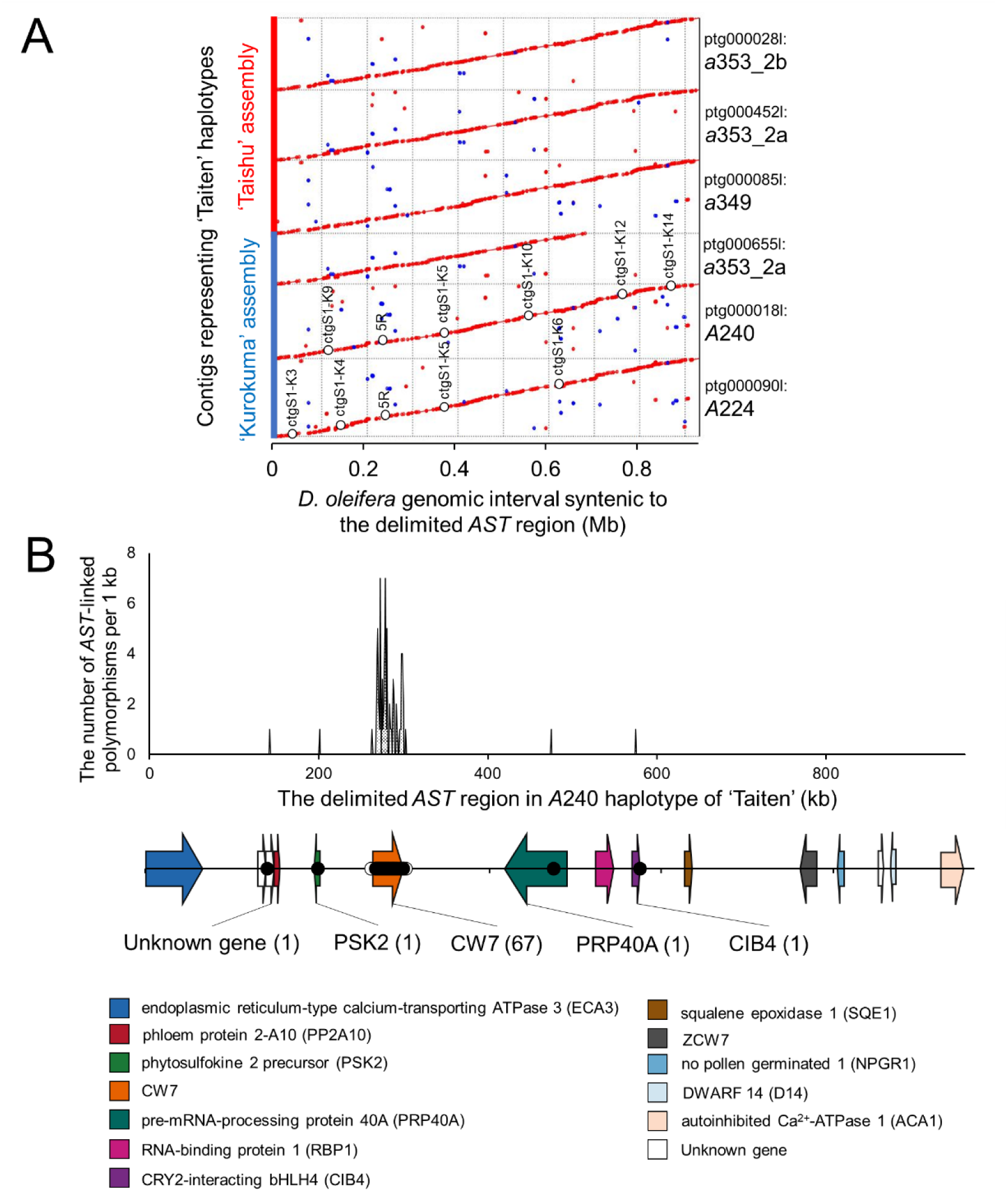
Identification of a single candidate gene at the *AST* locus by haplotype-resolved genome assembly and polymorphism analysis. (A) Synteny analysis of the delimited *AST* region between the corresponding region in the diploid *D. oleifera* genome^9^ and haplotype-resolved sequences of hexaploid *D. kaki* ‘Taiten’. Among six distinct haplotypes in ‘Taiten’, five haplotypes (*A*224, *A*240, *a*349, *a*353_2a, *a*353_2b) were successfully assembled, whereas *a*355 was not recovered. No detectable polymorphisms were found between *a*353_2b and *a*355 (fig. S3). Polymorphisms used for the Indel-3 marker (5R)^13^ and the *A*224- and *A*240-specific markers^15^ were detected in the corresponding *A*224 and *A*240 contigs. (B) Distribution of *AST*-linked polymorphisms detected in the delimited *AST* region of the *A*240 haplotype sequence. Arrows indicate coding regions and their transcriptional orientation. Numbers in parenthesis indicate the number of *AST*-linked polymorphisms within each gene region (from 5’UTR to 3’ UTR). Gene annotations are indicated by different colors shown in the bottom. The concentrated accumulation of *AST*-linked polymorphisms in a single gene annotated as CW7 (AT1G59520) indicates this gene as the strongest candidate at the *AST* locus.

Using the haplotype-resolved sequences as references, we aimed to identify polymorphisms that cosegregate with the presence of dominant *AST* allele(s) in hexaploid *D. kaki*. Whole genome sequencing data were obtained from 11 cultivars and from the 652-line breeding population (‘Fuyu’ x ‘Yoshidagosho’). The paternal parent of the 652-line population, ‘Yoshidagosho’, carries the *A*353 haplotype at the *AST* locus, which retains dominant *AST* function despite carrying the Indel-3 insertion typically associated with recessive *ast* haplotypes^20^. Because all reported *ast* haplotypes carry Indel-3 insertion, *A*353 is considered closely related to the *ast* haplotypes while retaining *AST* function (Supplementary Fig. 4). Therefore, sequence information derived from the 652-line population is highly informative for further refining candidate polymorphisms associated with PCNA trait.

A mapping-based variant calling approach, using the *A*224 and *A*240 haplotype sequences as a reference, identified polymorphisms within the delimited *AST* region that were heterozygous in genotypes carrying functional *AST* haplotype(s) and homozygous in genotypes lacking them (Supplementary Fig. 5). Gene positions in this region were annotated by full-length transcript (Iso-seq) mapping (Fig. 1B, Supplementary Table 5). Strikingly, candidate polymorphisms were strongly concentrated in a genomic region that harbors a single gene homologous to *CW7*, originally described in *Arabidopsis* (AT1G59520)^21^, although its biological function has not been characterized. Most polymorphisms were mapped to intron regions of this gene on the *A*240 reference (Supplementary Fig. 6). On the *A*224 reference, polymorphisms were more broadly distributed across the interval, yet 42 of 146 variants also concentrated within this gene (Supplementary Fig. 7). A few other genes detected using the *A*240 reference containing candidate polymorphisms showed no differences between PCNA and non-PCNA types in expression levels, and the detected polymorphisms do not influence nonsynonymous substitution or the gene splicing (Supplementary Fig. 8A, Supplementary Fig. 9, Supplementary Fig. 10). Therefore, the homolog of *CW7* was selected for further investigation as the sole strong candidate underlying the *AST* locus, and we designated this gene as *TANNIN* (*TAN*).

<u>Complex transcript variants in *DkTAN* caused by two mutations defining distinct *ast* alleles</u> Gene expression levels of *DkTAN* were not associated with the PCNA phenotype (Supplementary Fig. 8B). However, PCR analysis failed to detect cDNA encoding full-length DkTAN protein in PCNA samples, suggesting that transcript structure, rather than total expression level, may be associated with *DkTAN* dysfunction. We therefore examined whether *DkTAN* dysfunction is a consistent feature of PCNA individuals by amplicon sequencing of full-length cDNA from immature fruit. We first analyzed ‘Taiten’ (non-PCNA; *AAaaaa*) and ‘Kanshu’ (PCNA; *aaaaaa*). Both cultivars produced multiple transcript variants predicted to encode truncated proteins. In ‘Taiten’, transcripts encoding full-length functional proteins (389–392 aa) accounted for ∼30% of total *DkTAN* transcripts. In contrast, ‘Kanshu’ produced almost no functional transcripts (Fig. 2A; 1.0 %).

**Figure 2.**
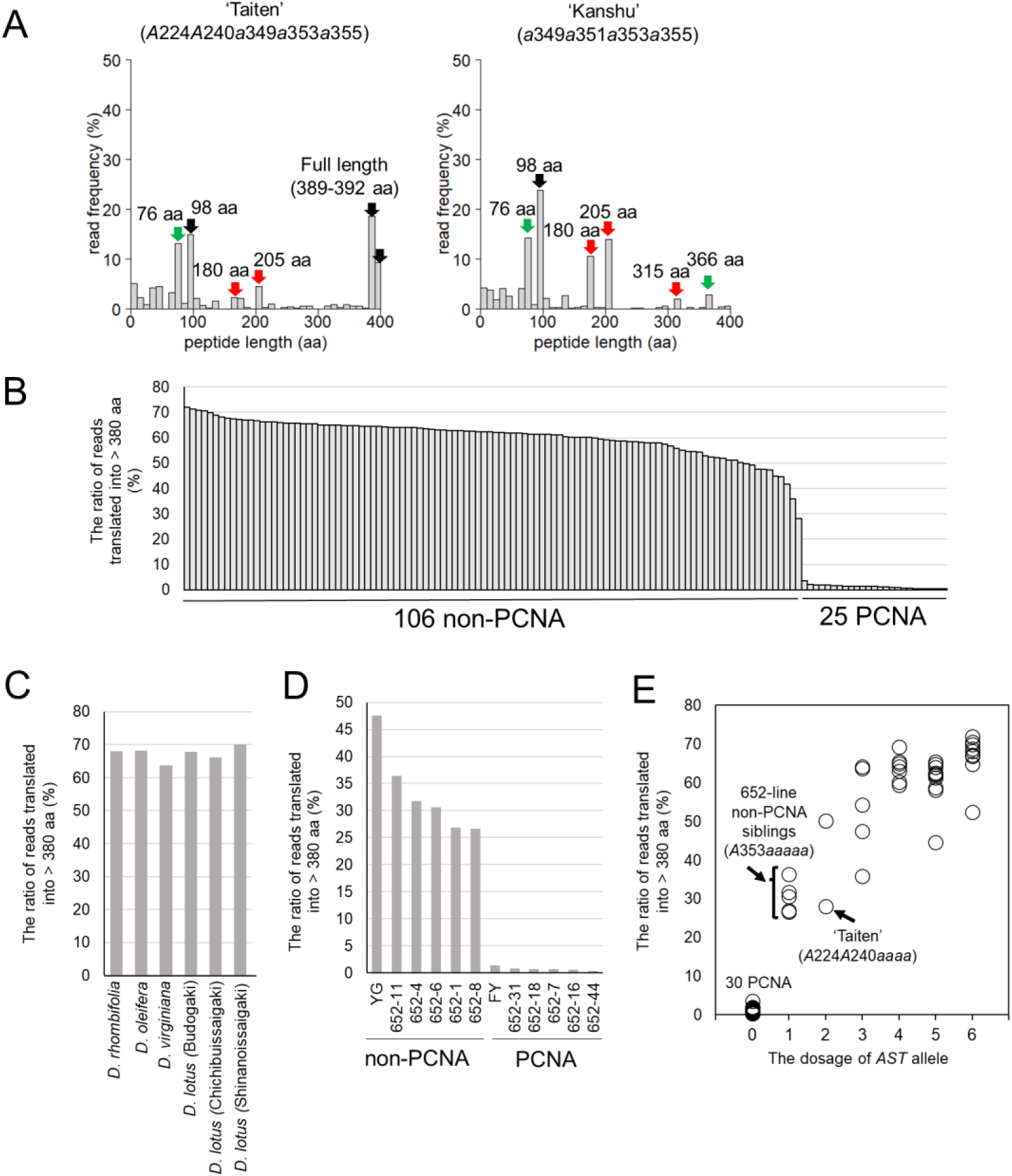
Abnormal *DkTAN* transcription is strongly associated with PCNA and non-PCNA genotypes. (A) Distribution of read frequencies obtained from cDNA amplicon sequencing of ‘Taiten’ and ‘Kanshu’, classified by predicted peptide length. Functional full-length translations (389– 392 aa) were detected in ‘Taiten’, whereas they were detected only at trace level in ‘Kanshu’. Black, green and red arrows indicate dominant peptide length translated from *AST*, *a*349 and *a*353 (*a*355), respectively. (B) Ratio of *DkTAN* transcripts encoding full-length functional proteins, estimated from amplicon-seq data using 106 non-PCNA and 25 PCNA cultivars. cDNA sequences predicted to encode proteins longer than 380 aa were classified as functional transcripts. A clear difference in the abundance of functional transcripts was observed between comprehensive non-PCNA and PCNA cultivars. (C) Ratio of functional *TAN* transcripts in *Diospyros* relatives. In all graphs, cDNA reads predicted to encode proteins longer than 380 aa were classified as functional transcripts. (D) Ratio of functional *DkTAN* transcripts in the 652-population (‘Fuyu’ x ‘Yoshidagosho’). Ten sibling individuals, including both PCNA and non-PCNA progeny, together with their parental cultivars, were analyzed. YG, ‘Yoshidagosho’; FY, ‘Fuyu’. (E) Relationship between the ratio of functional *DkTAN* transcripts and dominant *AST* dosage. *AST* dosage information of cultivars was obtained from a previous report^22^. The non-PCNA siblings from the 652-line population analyzed in this study were genotyped as *A*353*aaaaa* based on the genotyping of the Indel-3 region^20^.

To determine whether functional *DkTAN* transcription corresponds with the PCNA/non-PCNA phenotypes, we conducted amplicon sequencing across a comprehensive set of persimmon cultivars and *Diospyros* relatives (Supplementary Table. 6). Functional transcripts were detected only at trace levels in all available 25 PCNA landraces and modern cultivars (mean ± SE, 1.2 ± 0.14 %), whereas at least 25% of reads encoded full-length functional proteins in 106 non-PCNA cultivars and six related species (mean ± SE, 60.7 ± 0.69 %; Fig. 2B, C). The same pattern was observed in a segregating population (Fig. 2D). Moreover, the proportion of reads translating into full-length *DkTAN* increased with the dosage of the dominant *AST* alleles^22^ (Fig. 2E). Taken together, these data strongly support that the presence or absence of functional *DkTAN* transcripts is tightly associated with PCNA and non-PCNA phenotypes.

Comparison of the haplotype-resolved *DkTAN* sequences with the *D. oleifera* reference genome revealed two large indels present in haplotypes *a*353_2a and *a*353_2b but absent in *a*349 (Supplementary Fig. 11A). These included a 1.5-kb deletion that removes exon 7 and a 17-kb insertion within an intron. However, all representative PCNA cultivars are heterozygous for the 1.5-kb deletion (Supplementary Fig. 11B), indicating that it cannot be the sole causal mutation of all *ast* exists in PCNA cultivars.

To clarify the allelic basis of *DkTAN* dysfunction, we further analyzed the full-length amplicon sequencing data from ‘Taiten’ and ‘Kanshu’. Sequence analysis showed that the 1.5-kb deletion generates 180 aa, 205 aa, and 315 aa proteins through exon structure disruption (Fig. 2A, Fig. 3). However, additional truncated variants (76 aa, 98 aa, 366 aa) were detected in ‘Taiten’ and ‘Kanshu’ but not explained by the 1.5-kb deletion, indicating the presence of other causal mutation(s) associated with a distinct *ast* haplotype. From this point, we identified an *a*349-specific SNP (C→G) located at the 5′ splice site of exon 3, which is predicted to induce alternative splicing and exon skipping which cause 76 aa and 366 aa variants. The association between the candidate polymorphisms and *a*349 was confirmed using siblings from an *a*349-segregating population. Exon 3 SNP cosegregated with the *a*349, whereas the exon 7 deletion was absent from the *a*349 (Supplementary Fig. 12A, B), consistent with the haplotype-resolved genomic sequences (Supplementary Fig. 11). The predicted translation was verified by full-length cDNA amplicon sequencing (Supplementary Fig. 12C). Nonetheless, the 98 aa transcript variant is generated via exon reduction at the 3′ splice site, demonstrating that this variant is a natural isoform. Together, these results indicate that at least two independent mutations, the exon 7 deletion and a splice-site-related mutation around exon 3, lead to loss-of-function *DkTAN* alleles, each associated with a different *ast* haplotype group represented by *a*353_2a/*a*353_2b/*a*355 and *a*349, respectively (Fig. 3). Moreover, the identification of two independent loss-of-function allelic series, provides strong genetic support that *DkTAN* is the causal gene underlying the *AST* locus, consistent with established principles for inferring gene– trait causality^23^.

**Figure 3.**
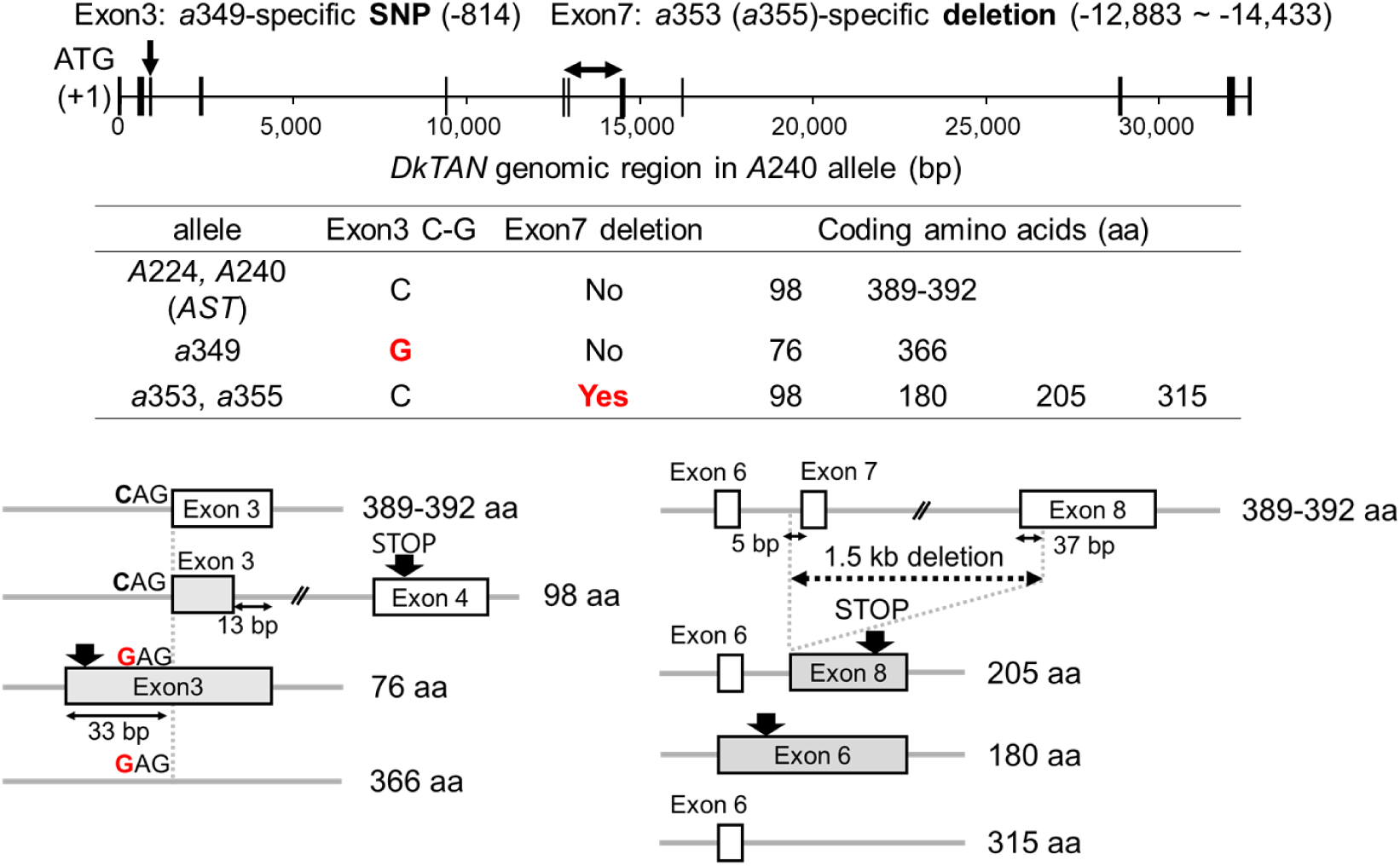
Complex abnormal *DkTAN* transcription caused by two independent mutations defining distinct *ast* haplotype groups. Schematic representation of abnormal transcripts caused by the two genomic mutations: the *a*349-specific SNP near exon 3 and the *a*353/*a*355-specific deletion spanning exon 7 and part of exon 8. The SNP at the 5′ splice site of exon 3 causes abnormal splicing variants, including shifts of the splice junction and exon 3 skipping. The 1.5-kb deletion including exon 7 and the first portion of exon 8 results in skipping of the corresponding exon regions. These abnormal transcripts generate premature stop codons and multiple truncated protein products. In addition, a transcript variant encoding a 98-aa peptide, caused by shortening of the 3′ region of exon 3, was detected independently of these two mutations. Deletion length and abnormal splicing model were inferred from haplotype sequence of ‘Taiten’ and Iso-seq mapping.

### Long-range haplotype-phased analysis for genotyping the two causal mutations

Although we identified two candidate causal polymorphisms, genotyping them across the germplasm collection required haplotype-phased approach because each polymorphism appeared in the heterozygous state when analyzed individually (Supplementary Fig. 13). However, the two candidate causal polymorphisms are separated by approximately 15 kb, making their simultaneous phasing highly challenging with conventional approaches. To achieve this, we adapted a target-enrichment approach combining probe-based hybridization with long-read sequencing (Supplementary Fig. 14). A total of 21 PCNA and four non-PCNA cultivars were analyzed, and a high enrichment efficiency was achieved within the *DkTAN* region selected for probe design in all cultivars (Supplementary Table 7). Following de novo assembly, haplotype sequences covering the *DkTAN* region were successfully reconstructed for each cultivar. Phylogenetic analysis together with the ‘Taiten’ haplotype assembly revealed that these sequences could be classified into three major groups: *AST*, *a*349, and *a*353 (Fig. 4A). Examination of the phased haplotype sequences demonstrated that the SNP near exon 3 was exclusively associated with the *a*349 group, whereas the deletion spanning exon 7 was specifically found in the *a*355 group (Fig. 4B). Importantly, all *DkTAN* sequences identified in PCNA cultivars belonged to either the *a*349 or *a*353 group. These phased genotyping results demonstrated that PCNA cultivars carries either of the two causal mutations across six *DkTAN* alleles, collectively accounting for the loss of functional *DkTAN* copies. This strongly supports the conclusion that these mutations are responsible for the functional deficiency of *DkTAN*.

**Figure 4.**
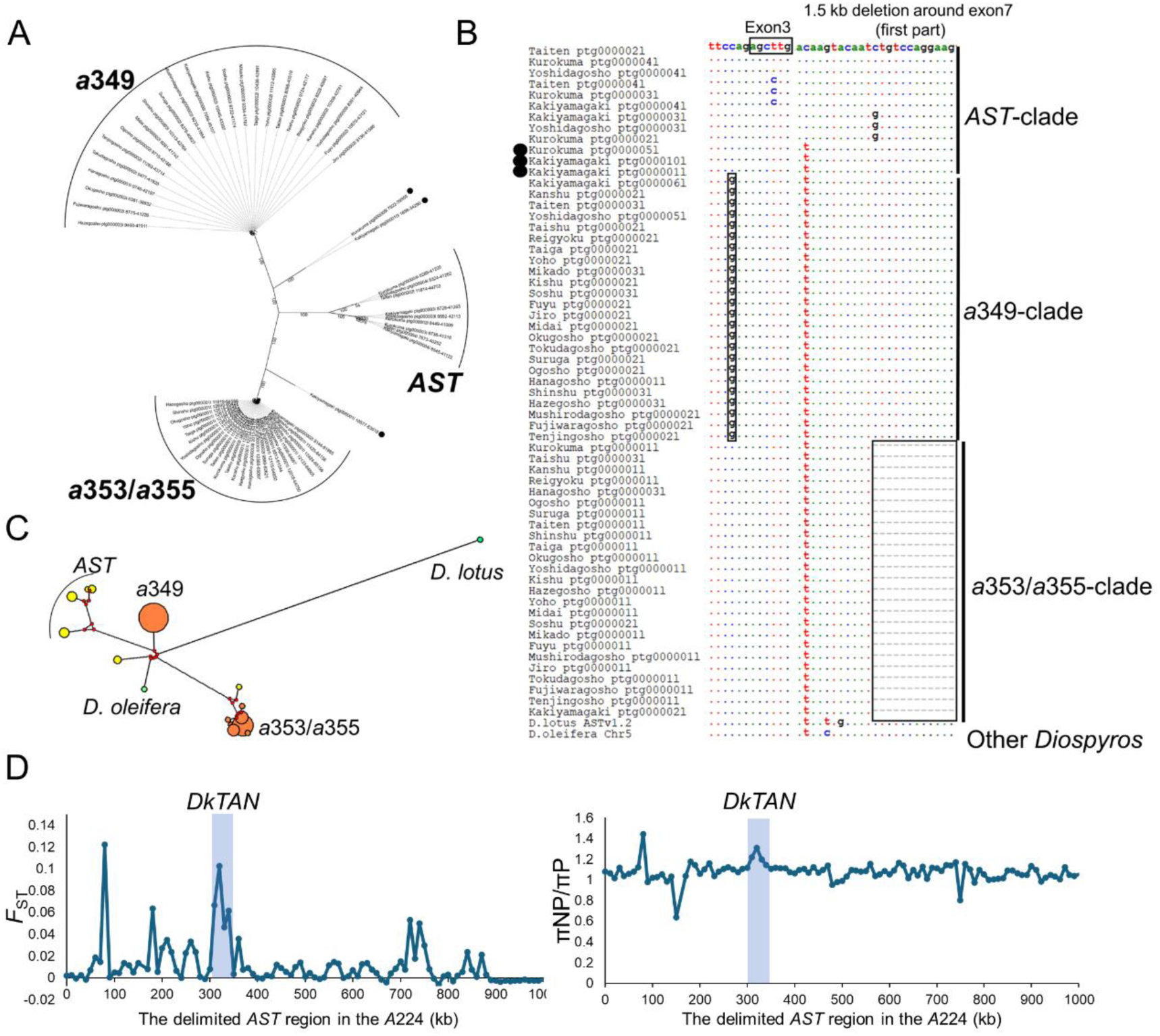
Sequence diversity of the *DkTAN* region revealed by long-read target-enrichment analysis in PCNA and non-PCNA cultivars. (A) Phylogenetic analysis of assembled haplotype-resolved sequences spanning the *DkTAN* genomic region. The tree topology suggests that *ast* haplotypes can be classified into two independent clades, *a*349 and *a*353/*a*355. In contrast, presumed functional *TAN* haplotype sequences from non-PCNA cultivars formed a distinct *AST* clade. Importantly, all sequences obtained from PCNA cultivars were classified into either the *a*349 or *a*353/*a*355 clade. (B) Phased genotyping of the two causal mutations using the assembled haplotype-resolved sequences. Sequence classification was based on the phylogenetic analysis shown in (A). All sequences belonging to the *a*349 clade possessed only the SNP near exon 3, whereas all sequences belonging to the *a*353/*a*355 clade possessed only the 1.5-kb deletion encompassing exon 7. *DkTAN* sequences from non-PCNA deviated from the *AST* clade (`●`) and those from other *Diospyros* species did not have either mutation. (C) Haplotype network analysis of assembled *DkTAN* sequences. Circle diameter represents the number of haplotypes included in each group, and line length indicates the sequence dissimilarity between connected haplotype groups. Yellow, orange, and green circles indicate *AST*, *ast*, and other *Diospyros* haplotype groups, respectively. Red dots represent median vectors, which indicate intermediate haplotypes inferred in the network analysis. (D) Population genetic analysis in the *AST* locus. Nucleotide diversity (π) and *F*_ST_ were calculated in 10 kb window using VCFtools based on DNA-seq mapping data from the cultivars used for polymorphism analysis. Variant calling was performed using VarScan, and the genotypes of the detected polymorphisms were refined using custom Python scripts based on alternative-allele read frequency thresholds optimized for the hexaploid genome (see Materials and Methods). Although a clear *F*_ST_ peak was detected around the *DkTAN* (Chr5: 315,216–348,804), π showed only a weak local increase, with no evidence of reduced nucleotide diversity in the PCNA group across the *DkTAN* region. The genomic region containing *DkTAN* is highlighted in blue. πP and πNP represent nucleotide diversity within the PCNA and non-PCNA populations, respectively.

### Independent *DkTAN* mutations underlying PCNA while maintaining haplotype diversity of the *AST* locus

The clear phylogenetic separation between the *a*349 and *a*353 groups suggests that these *ast* haplotypes emerged independently (Fig. 4A). Notably, some *DkTAN* haplotypes from non-PCNA cultivars were closely related to the *a*349 or *a*355 groups despite lacking causal mutations, suggesting the presence of intermediate haplotypes in non-PCNA. Haplotype network analysis further revealed that *TAN* homolog from *D. oleifera* were more closely related to *DkTAN* than those from *D. lotus*, consistent with previous reports proposing *D. oleifera* as one of the closest diploid ancestors of *D. kaki*^24^ (Fig. 4C). In addition, it was found that *a*349 haplotype group was closer to *D. lotus* and *D. oleifera* sequences, suggesting earlier emergence of *a*349 from *AST* than *a*353. Together, these findings suggest that the PCNA trait became established through the accumulation of multiple independently arisen *ast* haplotypes within the hexaploid genome, while diverse functional *AST* haplotypes were retained in non-PCNA cultivars (Supplementary Fig. 15). Consistent with this scenario, no evidence of a selective sweep in PCNA cultivars was detected across the *AST* locus, including the *DkTAN* region (Fig. 4D), suggesting that sequence diversity has been maintained among *ast* haplotypes. Thus, the establishment of PCNA appears to have resulted from accumulation of independent *ast* alleles rather than fixation of a single mutational event.

## Discussion

The massive accumulation of PAs in persimmon fruit represents one of the most extreme examples of specialized metabolite accumulation among fruit crops. Cultivar differences related to pollination response and fruit astringency have been recognized for more than a century and remain central to persimmon classification and breeding^25^. Although the *AST* locus has long been recognized as the major determinant of the PCNA phenotype, the underlying gene remained unknown because of the highly heterozygous hexaploid genome of persimmon. Here, long-read-assisted forward genetics identified *DkTAN* as the causal gene underlying *AST* and revealed that PCNA arose through at least two independently derived loss-of-function haplotypes (Fig. 1, Fig. 2, Fig. 3). Because PCNA is a recessive trait in a hexaploid species, breeding of PCNA cultivars has relied on repeated use of a limited number of elite founders, leading to a narrow genetic background and inbreeding depression-like constraints ^12^. In contrast, our results show that the *AST* locus itself retains haplotype diversity through multiple independent *DkTAN* loss-of-function alleles. This finding suggests that the unfavorable genetic constraints associated with PCNA breeding may not be explained solely by reduced diversity at the *AST* locus, and raises the possibility that loss of TAN function itself, or genetic factors inherent to PCNA founder backgrounds, contributes to these constraints. Thus, identification of *DkTAN* provides both an evolutionary framework for the recent establishment of PCNA cultivars and a molecular basis for more precise marker-assisted breeding (Fig. 4)^12, 20^.

Given the distinctive pattern of PA accumulation in persimmon compared with other plant species and its recently reported unique cellular sequestration system^26^, TAN may play a central role in the exceptionally high accumulation of PA specific to persimmon fruit. As TAN has not been characterized in any plant species, further investigation of its molecular function may provide new insight into both the mechanism underlying natural astringency loss in persimmon and large-scale sequestration of specialized metabolites in plants.

## Materials and Methods

### Plant materials and DNA sequencing

Young leaves were collected from the persimmon cultivar ‘Taiten’ grown at the Grape and Persimmon Research Station of the Institute of Fruit Tree and Tea Science, National Agriculture and Food Research Organization (NIFTS; Higashi-Hiroshima, Japan). Nuclei were isolated from leaf tissues following a modified version of the previous protocol^27^. High-molecular-weight DNA was subsequently extracted using the Nanobind DNA Extraction Kit (Circulomics), and DNA fragments shorter than 5 kb were removed using the Short Read Eliminator Kit (Circulomics). A total of 70 μg of high-molecular-weight DNA was used for library preparation and sequenced across seven SMRT Cells on the PacBio Sequel II platform. HiFi reads (≥Q20) were generated from the resulting subreads using packages available in pbbioconda (https://github.com/PacificBiosciences/pbbioconda). In total, approximately 185 Gb of HiFi reads were obtained from ‘Taiten’, corresponding to approximately 43× coverage of the hexaploid persimmon genome.

For polymorphism analysis, 13 cultivars, including the parental cultivars of ‘Taiten’ (‘Kurokuma’ and ‘Taishu’), grown at the Kyoto University Experimental Farm and NIFTS, were used. In addition, ten progeny individuals homozygous for Indel-3 from the ‘Fuyu’ × ‘Yoshidagosho’ population (652-line), generated and maintained at NIFTS, were included in the analysis. The ten progeny individuals were selected based on sensory evaluation data obtained at NIFTS so that the selected set included five non-PCNA and five PCNA individuals. Genomic DNA was extracted from young leaves using the CTAB method^28^. Short-read sequencing data from the cultivar panel and the 652-line population were generated using the Illumina platform. The resulting reads were subjected to quality filtering before downstream analyses. The sequencing results and accession numbers were summarized in Supplementary Table 1.

### Haplotype-resolved assembly of ‘Taiten’ genome

Trio-binning was performed using a modified version of the publicly available pipeline (https://github.com/esrice/trio_binning)^18, 29^. Short-read sequencing data from the parental cultivars ‘Kurokuma’ and ‘Taishu’ were decomposed into k-mers using the find-unique-kmers function, and parent-specific k-mers were identified. Independent analyses were conducted using multiple k-mer lengths (k = 17, 19, 21, 23, 25, 27, 31, and 37). To evaluate the optimal k-mer length, 254 HiFi reads spanning the 5R region^13^ were manually assigned to their corresponding haplotypes based on sequence polymorphisms. Using the classify_by_kmers function, HiFi reads containing the 5R region were classified into the ‘Kurokuma’ haplotype group (hapA), the ‘Taishu’ haplotype group (hapB), or an unclassified group (hapU). Reads were assigned to hapA or hapB when they contained at least 10 parent-specific k-mers and more than 90% of the parent-specific k-mers were derived from a single parent (Supplementary Fig. 1). For each k-mer length, the numbers of correctly classified reads in hapA and hapB (a and b, respectively) were summed, and the partition accuracy was calculated as (a + b)/254 (Supplementary Table 2).

Because k = 25 and k = 31 showed the highest partition accuracy (see main text), these k-mer lengths were subsequently used for genome-wide classification of HiFi reads. The classified read sets corresponding to the ‘Kurokuma’ haplotype group (hapA + hapU) and the ‘Taishu’ haplotype group (hapB + hapU) were independently assembled using hifiasm^30^ with the parameters --n-hap 3 and -l 0. Primary contigs corresponding to the *AST* locus region were then identified. The haplotype origin of each contig was determined using marker sequences located within the 5R region. Finally, sequence similarity between the assembled contigs and the *AST* locus region of the *D. oleifera* reference genome^19^ was assessed using NUCmer from the MUMmer package v3.23^31^.

### Read mapping, variant calling, and identification of *AST*-linked polymorphisms

The workflow used for the identification of *AST*-linked polymorphisms is summarized in Supplementary Fig. 4. Short-read DNA sequencing data obtained from the cultivars and populations described above were mapped independently to the *A*224 and *A*240 haplotype sequences assembled from ‘Taiten’, focusing on the delimited *AST* region^15^. Read mapping was performed using BWA-MEM v0.7.17^32^. BAM files were generated using SAMtools view ^33^. For the 652-line population, BAM files from five PCNA individuals and five non-PCNA individuals were merged separately. Variant calling detecting single nucleotide polymorphisms (SNPs) and short insertions/deletions (short indels) was performed using SAMtools mpileup. Variant call datasets from the cultivar panel and the 652-line population were subsequently merged using custom Python scripts.

To eliminate false-positive variants originating from repetitive regions with excessive read accumulation, coverage filtering was applied prior to variant screening. For ‘Kanshu’, ‘Taishu’, ‘Taiten’, and ‘Kurokuma’, only sites with read depths between 20× and 300× were retained. For all other cultivars, a threshold of 50×–300× was applied. Sites outside these ranges were excluded from further analyses. Among the retained variants, the candidate *AST*-linked polymorphisms were defined as sites that were homozygous in all PCNA cultivars (major allele ratio = 100%) and heterozygous in non-PCNA cultivars (minor allele ratio ≥ 5%). To further validate linkage with the phenotype, variant calls from the 652-line population were extracted only at the candidate *AST*-linked sites detected in the diversity panel. Variants were retained when the major allele ratio was 100% in both ‘Fuyu’ and the pooled PCNA progeny, while the minor allele ratio was at least 15% in both ‘Yoshidagosho’ and the pooled non-PCNA progeny.

Finally, variants were further filtered to retain only those for which PCNA cultivars were homozygous for the alternative allele, defined as an alternative allele frequency of at least 99% relative to the reference sequence. These variants were designated as *AST*-linked polymorphisms. The distribution of identified *AST*-linked polymorphisms across the delimited *AST* region in the *A*224 and *A*240 haplotype sequences was visualized using 1-kb genomic bins.

### Iso-Seq library preparation, sequencing, and analysis

Mesocarp tissues were collected from ‘Taiten’ fruits harvested in June, July, and August 2016. Total RNA was extracted using the hot borate method^34^. RNA quality was assessed using a Qubit 4 Fluorometer (Invitrogen), and only samples with an RNA IQ value greater than 9 were used for library preparation. Full-length transcript sequencing (Iso-Seq) was performed on the PacBio Sequel II platform. Raw sequence data were processed using SMRT Link (PacBio) to generate high-quality Iso-Seq reads for each sampling date.

Iso-Seq datasets obtained from the three developmental stages were merged and mapped to the delimited *AST* regions in the *A*224 and *A*240 haplotype sequences of ‘Taiten’ using minimap2^35^. Mapped reads were retrieved using SAMtools view with the -F 2048 option to remove supplementary alignments. To eliminate spurious alignments, reads were further filtered by local BLAST analysis, retaining only those for which more than 50% of the transcript sequence was aligned to the target region. The filtered reads were subsequently remapped to the delimited *AST* regions in the *A*224 and *A*240 haplotypes using minimap2. Mapping results were visualized using Integrative Genomics Viewer (IGV)^36^, and the positions of expressed genes within the delimited *AST* region were manually annotated based on transcript alignments.

### Amplicon sequencing

A comprehensive germplasm panel consisting of 131 persimmon cultivars, six related *Diospyros* accessions, and ten progeny individuals from the 652-line population were used for amplicon sequencing analysis (Supplementary Table 6). Fruits were collected between late June and early July, and total RNA was extracted from mesocarp tissues using the ISOSPIN Plant RNA Kit (Nippon Gene) according to the manufacturer’s instructions with Assist Buffer for ISOSPIN Plant RNA (Nippon gene). First-strand cDNA was synthesized from oligo (dT)-purified RNA (total 1 µg) using SuperScript III Reverse Transcriptase (Invitrogen). The full-length coding sequence of the candidate gene, including the start and stop codons, was amplified by a first-round PCR using KOD One® PCR Master Mix (TOYOBO). Sample-specific index sequences were subsequently added in a second-round PCR. Primer sequences are listed in Supplementary Table 8. To minimize amplification bias, the numbers of PCR cycles were limited to 25 and 10 cycles for the first and second PCRs, respectively. The resulting libraries were subjected to full-length amplicon sequencing on the PacBio Sequel platform. Following demultiplexing based on index sequences, cDNA sequences were translated into amino acid sequences using gotranseq (https://github.com/feliixx/gotranseq). Translated sequences were further processed using custom Python scripts to remove amino acid sequences downstream of premature stop codons. Amino acid sequences encoding full-length proteins without premature stop codons were classified as functional transcripts.

### Target enrichment sequencing and haplotype analysis

The workflow is summarized in Supplementary Fig. 15. Young leaves were collected from four non-PCNA and 21 PCNA cultivars grown at the Kyoto University Experimental Farm. Approximately 5 g of leaf tissue was used for genomic DNA extraction following the CTAB method. Purified DNA was mechanically sheared using a Megaruptor^®^ system (Diagenode) according to the manufacturer’s instructions. Fragment size distributions were evaluated for representative samples using a Femto Pulse System (Agilent), confirming that most DNA fragments were concentrated around 7–8 kb. Sequencing libraries were prepared from the sheared DNA using the GenNext^®^ NGS Library Prep Kit (TOYOBO), followed by ligation of TruSeq Combinatorial Dual Index adapters (Illumina). Target enrichment was performed using a custom probe panel, designed by Twist Bioscience, against the candidate gene region in the ‘Taishu’ genome (Chr5: 16446515-16509450)^37^. Capture enrichment was conducted according to the Long Read Library Preparation and Standard Hyb v2 Enrichment protocol (Twist Bioscience). Two independently prepared multiplex libraries were pooled and sequenced on the PacBio Revio platform.

Following sequencing, reads were demultiplexed according to sample-specific index sequences. HiFi reads longer than 5 kb were assembled de novo using hifiasm with the parameters -l 0 and --n-hap 6. Primary contigs were aligned to the ‘Taiten’ haplotype sequences (*A*240 and *a*353_2b) using minimap2, and contigs corresponding to the candidate gene region were extracted using custom Python scripts. Contigs longer than 20 kb for the *A*240 haplotype and longer than 40 kb for the *a*353_2b haplotype were retained and combined for subsequent analyses. The selected sequences were aligned using MAFFT v7^38^ with the --auto option. The resulting multiple sequence alignment was subjected to phylogenetic analysis using IQ-TREE 3^39^, with automatic model selection (-m MFP) and 1,000 ultrafast bootstrap replicates (-B 1000). Phylogenetic trees were visualized using the Interactive Tree of Life (iTOL; https://itol.embl.de/).

For haplotype network analysis, the MAFFT alignment was converted to NEXUS format and imported into DnaSP v6^40^ to define haplotype groups. The resulting data were exported in RDF format and analyzed using NETWORK v10.2 (Fluxus Technology Ltd., https://www.fluxus-engineering.com/index.htm). Haplotype networks were constructed using the median-joining algorithm.

### Sequence diversity analysis

BAM files generated for the polymorphism analysis from nine PCNA cultivars (‘Fuyu’, ‘Okugosho’, ‘Kanshu’, ‘Jiro’, ‘Hanagosho’, ‘Fujiwaragosho’, ‘Mikado’, ‘Midai’, and ‘Taishu’) and four non-PCNA cultivars (‘Taiten’, ‘Tenryubo’, ‘Kuromuka’, and ‘Yoshidagosho’) were used for variant calling with SAMtools mpileup and VarScan v2.3^41, 42^, using the ‘Taiten’ *A*224 haplotype sequence spanning the delimited *AST* region as the reference. In the resultant Variant Call Format (VCF) file, biallelic SNPs were initially filtered using VCFtools v0.1.17^43^ with the options (--minDP 20 --max-meanDP 200 --max-missing 0.8 --mac 4 --min-alleles 2 --max-alleles 2 --remove-indels). Because VarScan is not designed for hexaploid genotype calling, genotype calls were further refined using custom Python scripts based on allele frequencies. Genotypes with alternative-allele read frequencies below 0.3% were converted to homozygous reference calls, whereas those above 99.7% were converted to homozygous alternative calls. Genotypes with alternative-allele read frequencies between 0.3% and 5% or between 95% and 99.7% were masked as missing to reduce ambiguous low-frequency calls, and those between 5% and 95% were treated as heterozygous. Population genetic statistics, including *F*_ST_ and nucleotide diversity (π), were then calculated for the PCNA and non-PCNA populations using VCFtools in non-overlapping 10-kb windows.

## Data availability

All sequencing data generated in this study have been deposited in the DDBJ Sequence Read Archive under accession number PRJDB42921.

