## Supplementary Materials for "Accumulation of two distinct alleles causes astringency loss in cultivated polyploid persimmon"

### Supplementary Figures

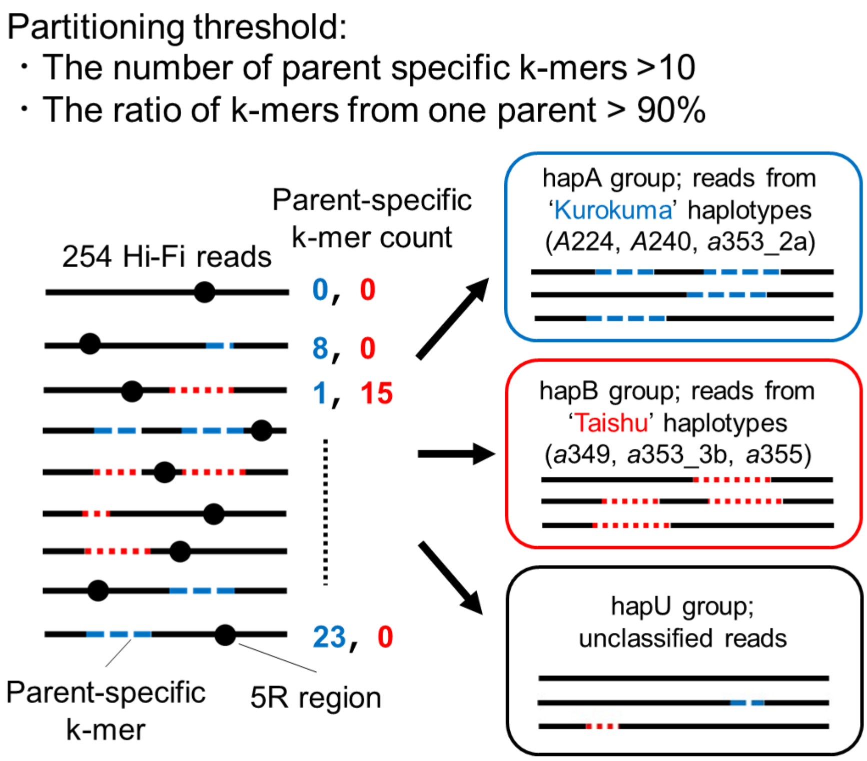


**Supplementary Fig. 1 Trio-binning approach for 'Taiten' assembly**

HiFi reads generated from 'Taiten' genomic DNA were classified into 'Kurokuma'-derived haplotype group (hapA) and 'Taishu'-derived haplotype group (hapB) based on thresholds of parent-specific k-mer counts. For partitioning test of each k-mer length, 254 HiFi reads possessing 5R region (*13*) were retrieved and partitioned.

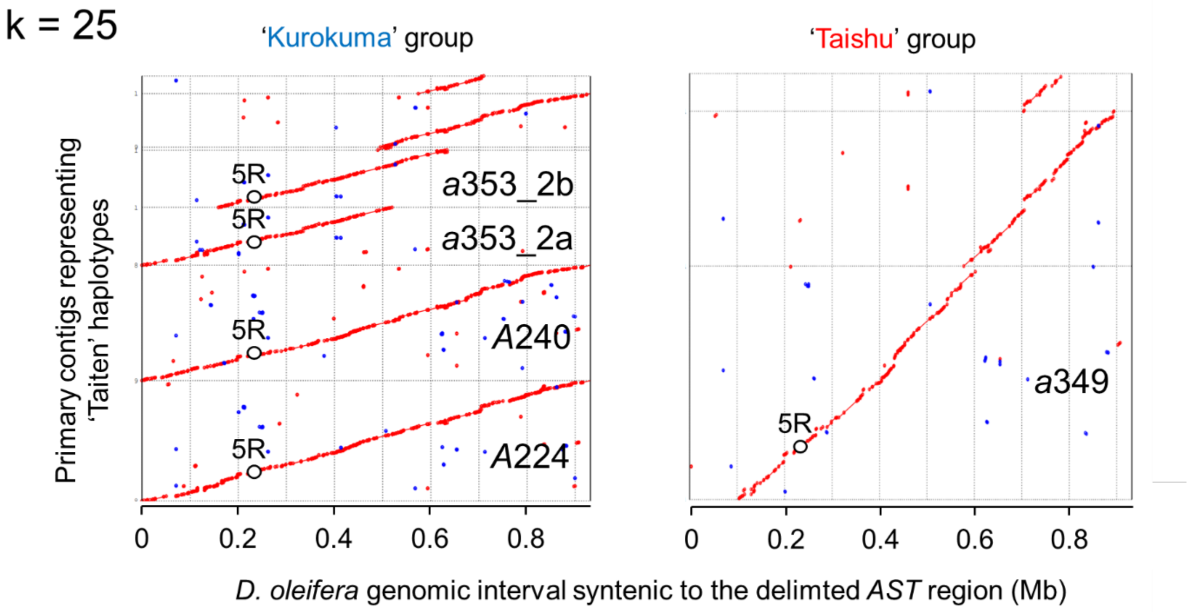


**Supplementary Fig. 2 Alignment of the delimited *AST* region in trio-binned assemblies generated from whole genome HiFi reads partitioning with k = 25**

Synteny analysis of the 'Kurokuma' and 'Taishu' assemblies are shown separately. 5R indicates the Indel-3-containing region used to distinguish *AST* and *ast* haplotypes.

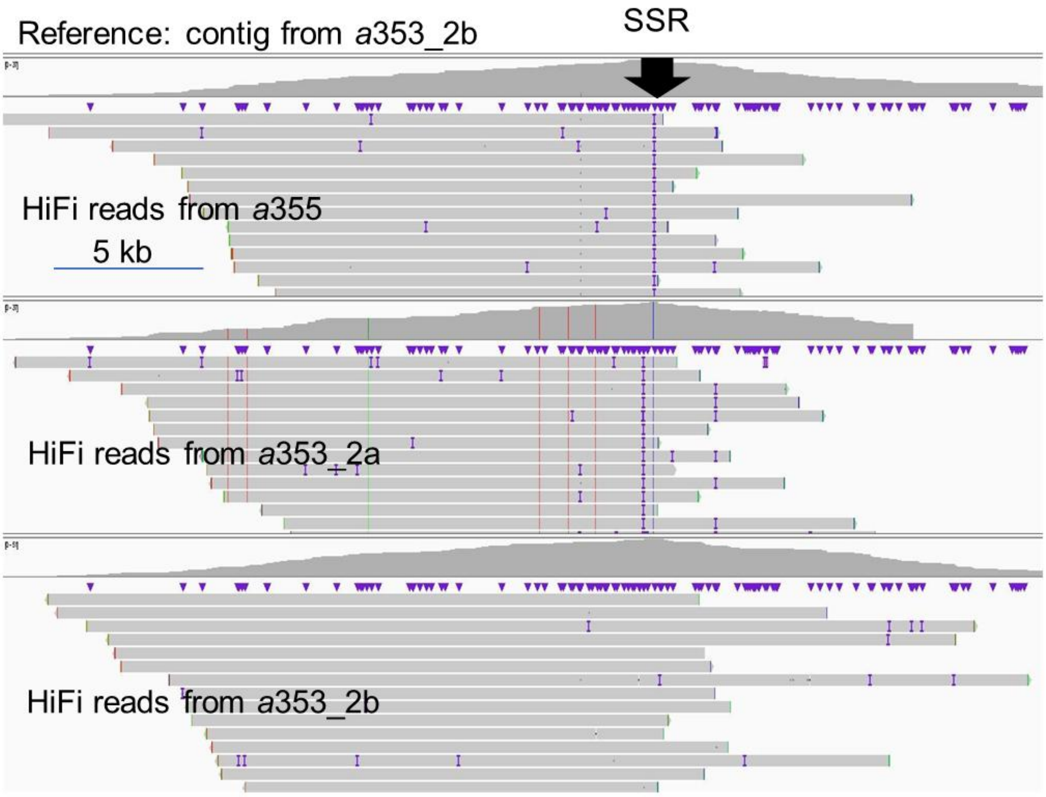


**Supplementary Fig. 3 Mapping of HiFi reads from *a*355, *a*353_2a and *a*353_2b using *a*353_2b as the reference sequence**

Mapping results were visualized using IGV. The results indicate that *a*355 has no detectable polymorphisms relative to *a*353_2b, except for a short sequence repeat (SSR) around Indel-3.

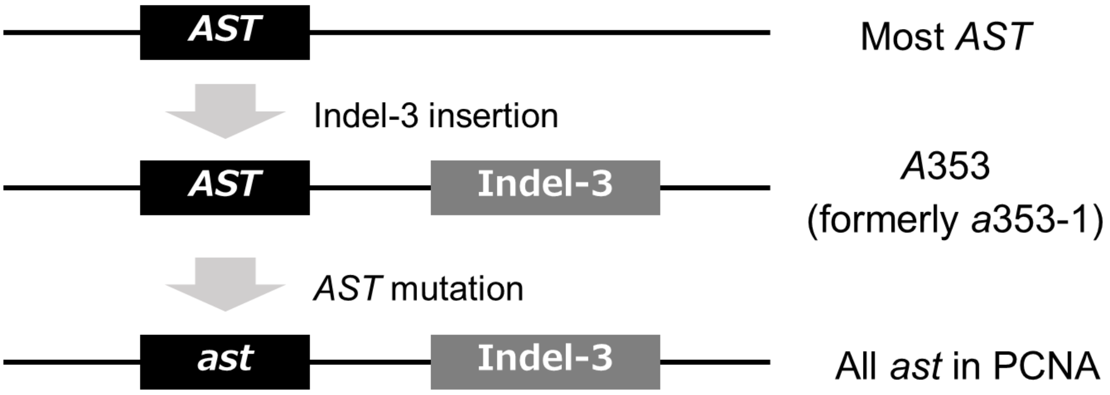


**Supplementary Fig. 4 Proposed scheme for the establishment of the *A*353 haplotype according to Indel-3 insertion and *AST* haplotype function**

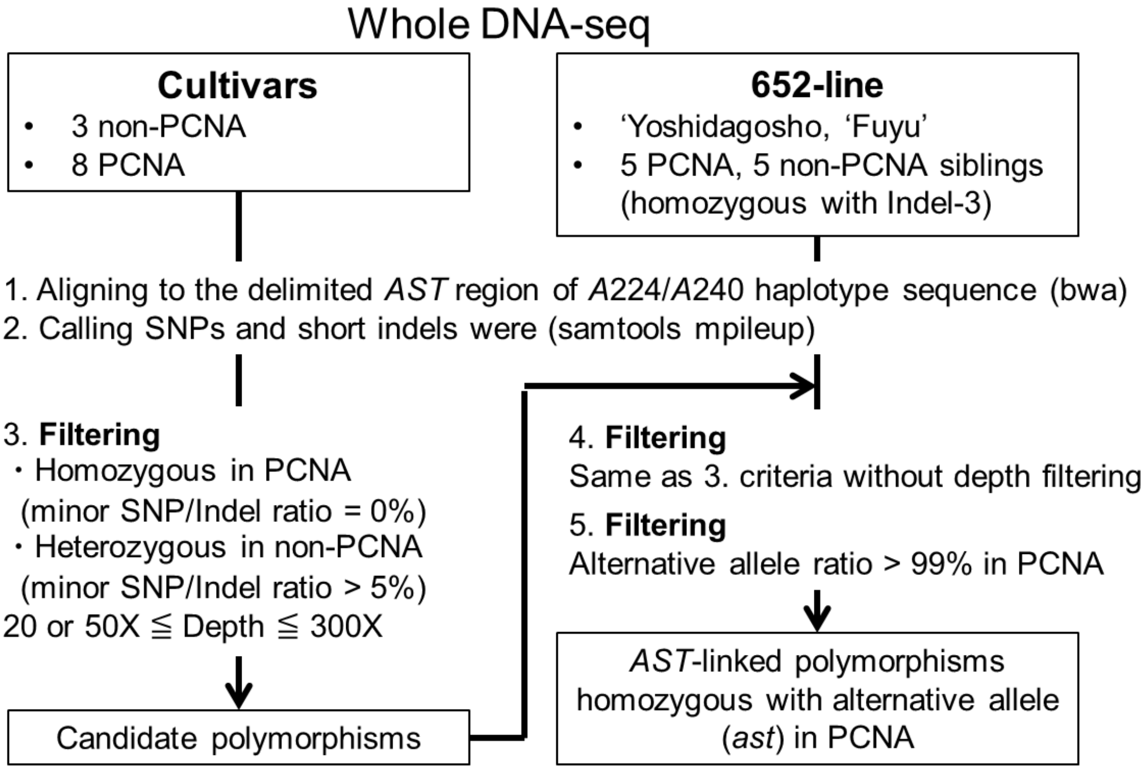


**Supplementary Fig. 5 Workflow for the *AST*-linked polymorphism detection**

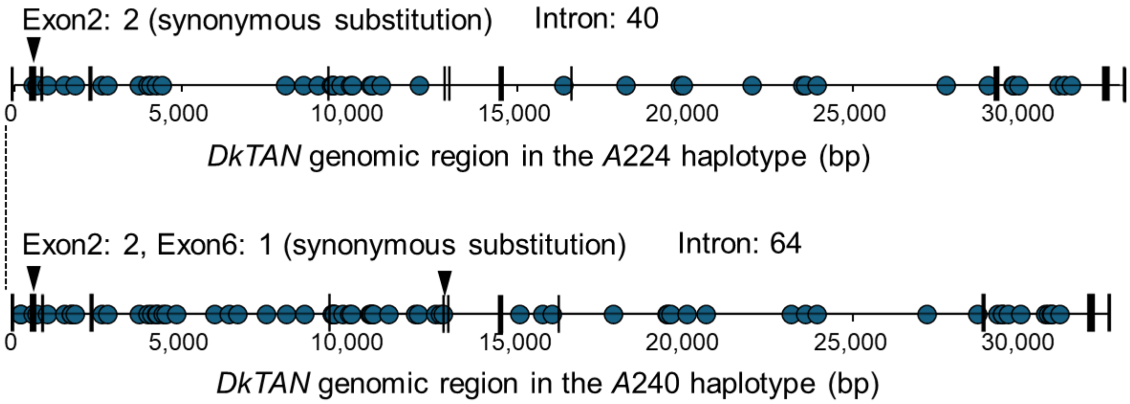


**Supplementary Fig. 6 *DkTAN* gene model and the distribution of *AST*-linked polymorphisms detected in the *A*224 and *A*240 haplotypes**

Blue dot and vertical line indicate the position of *AST*-linked polymorphisms and exon, respectively. Black arrow shows the position of polymorphisms in exon. All other polymorphisms were located in intron regions. A few polymorphisms were detected in exonic regions, but all were synonymous substitutions.

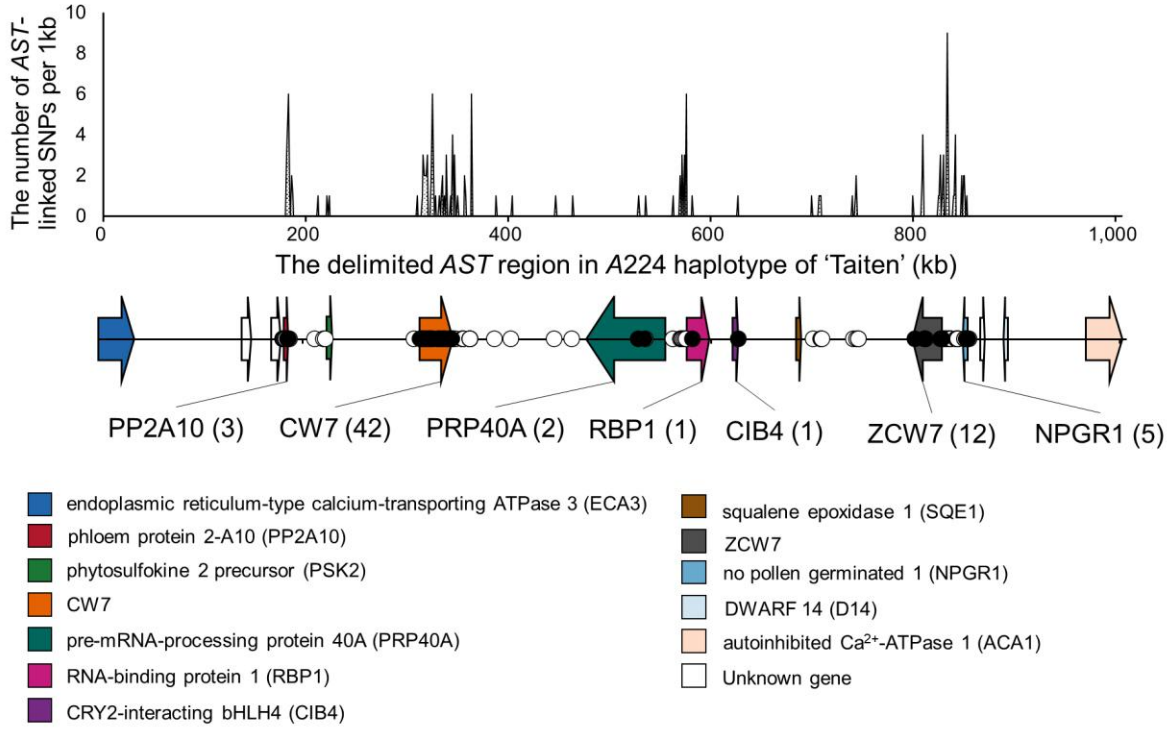


**Supplementary Fig. 7 Distribution of *AST*-linked polymorphisms detected in the delimited *AST* region of the *A*224 haplotype sequences**

The detail explanation follows caption in Figure 1B.

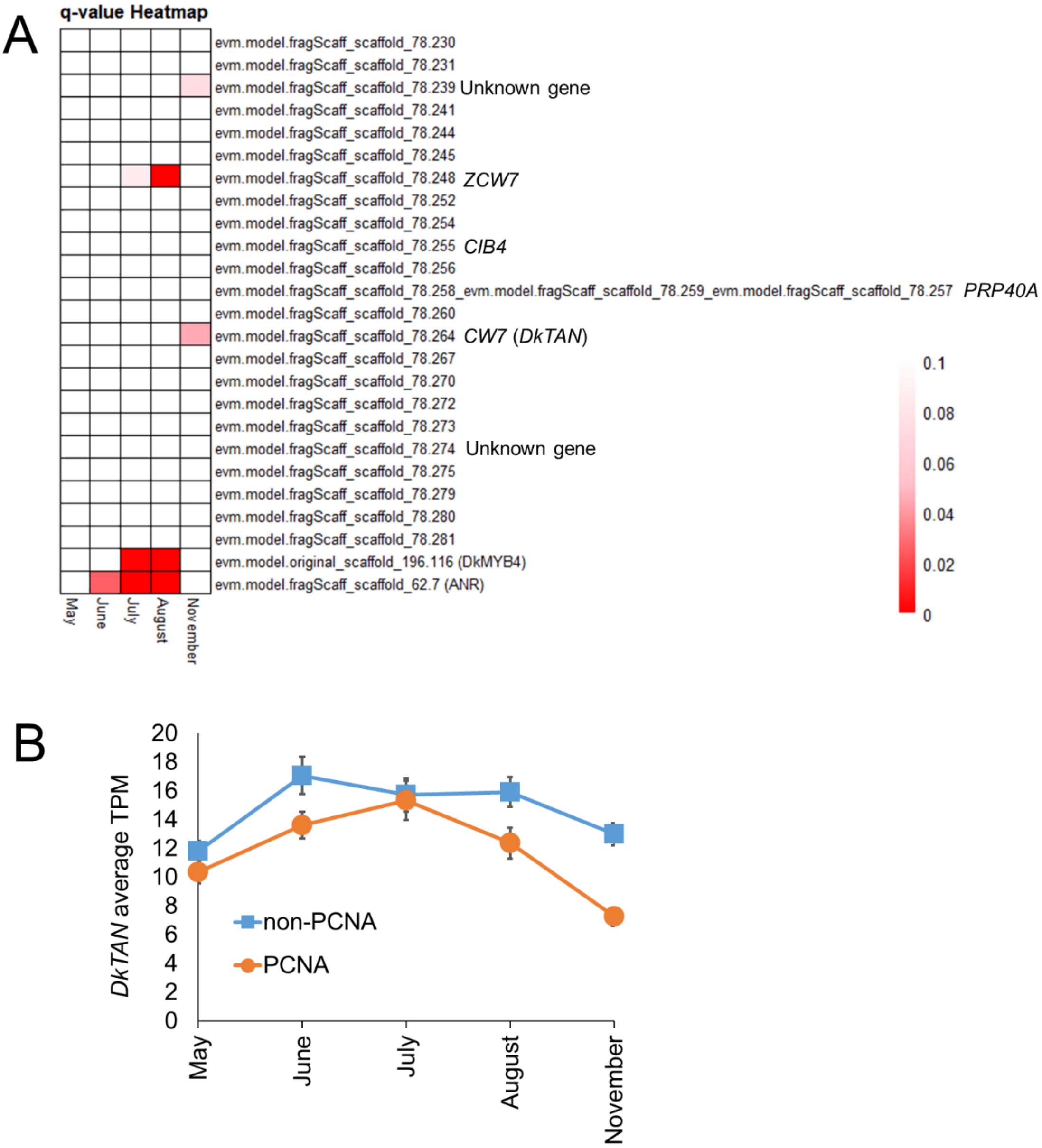


**Supplementary Fig. 8 Expression differences between PCNA and non-PCNA individuals for genes in the delimited *AST* region and PA-related genes**

(A) Heatmap showing differential expression level of genes. q-values were obtained from our previous study (*16*), in which differential expression gene (DEG) analysis was performed using an $F_1$ population derived from a cross between 'Taiten' (non-PCNA) and 'Kanshu' (PCNA). The analysis was based on fruit RNA-seq data collected across five developmental stages. As positive controls for expression differences associated with PA accumulation, q-values for *DkMYB4* and *ANR* were included because these genes have previously been reported to exhibit expression patterns consistent with PA accumulation differences between PCNA and non-PCNA (*8*). Among the genes located within the delimited *AST* region, none showed expression patterns corresponding to the PA accumulation phenotype. (B) Expression profile of *DkTAN* in PCNA and non-PCNA siblings. Expression data were retrieved from our previous study (*16*). Error bars indicate standard errors.

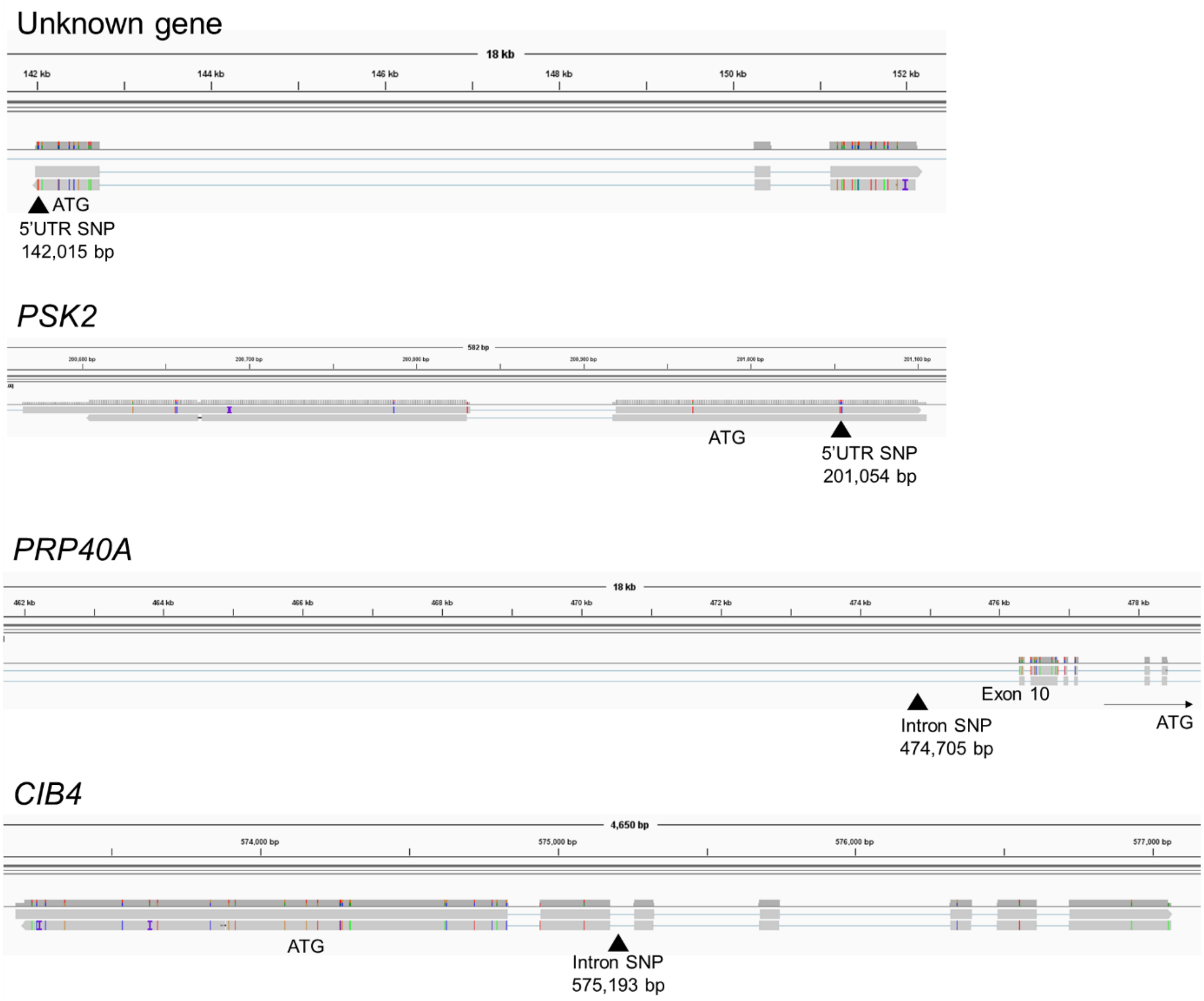


**Supplementary Fig. 9 Iso-Seq read mapping of genes other than *DkTAN* that contain *AST*-linked polymorphisms**

The contig representing the *A*240 haplotype was used as the reference sequence. In the IGV visualization, reads without colored mismatches indicate transcripts derived from the *AST* haplotype of 'Taiten', while reads with colored SNPs represent transcripts derived from *ast* haplotypes of 'Kanshu' or 515-118 (PCNA from 'Taiten' x 'Kanshu').

### CIB4

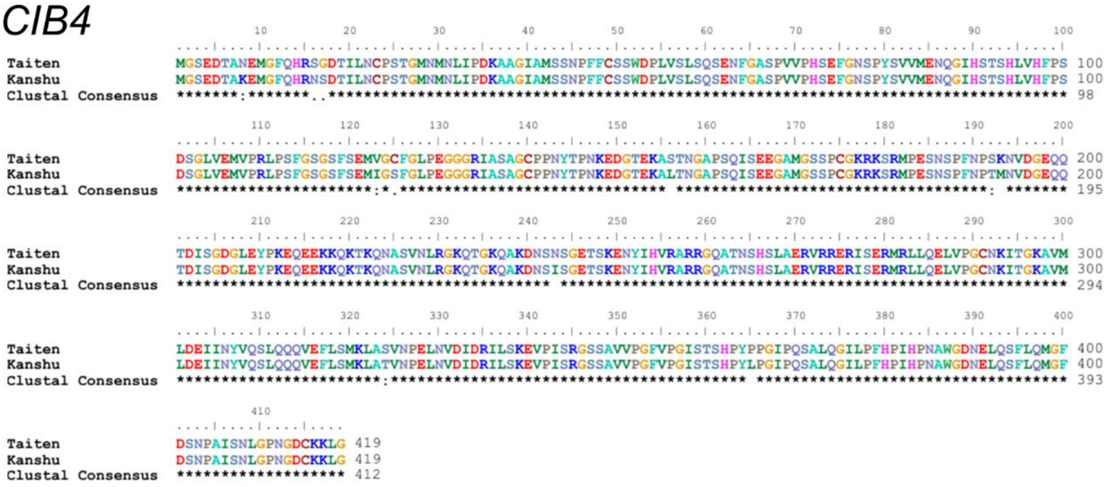

### PRP40A

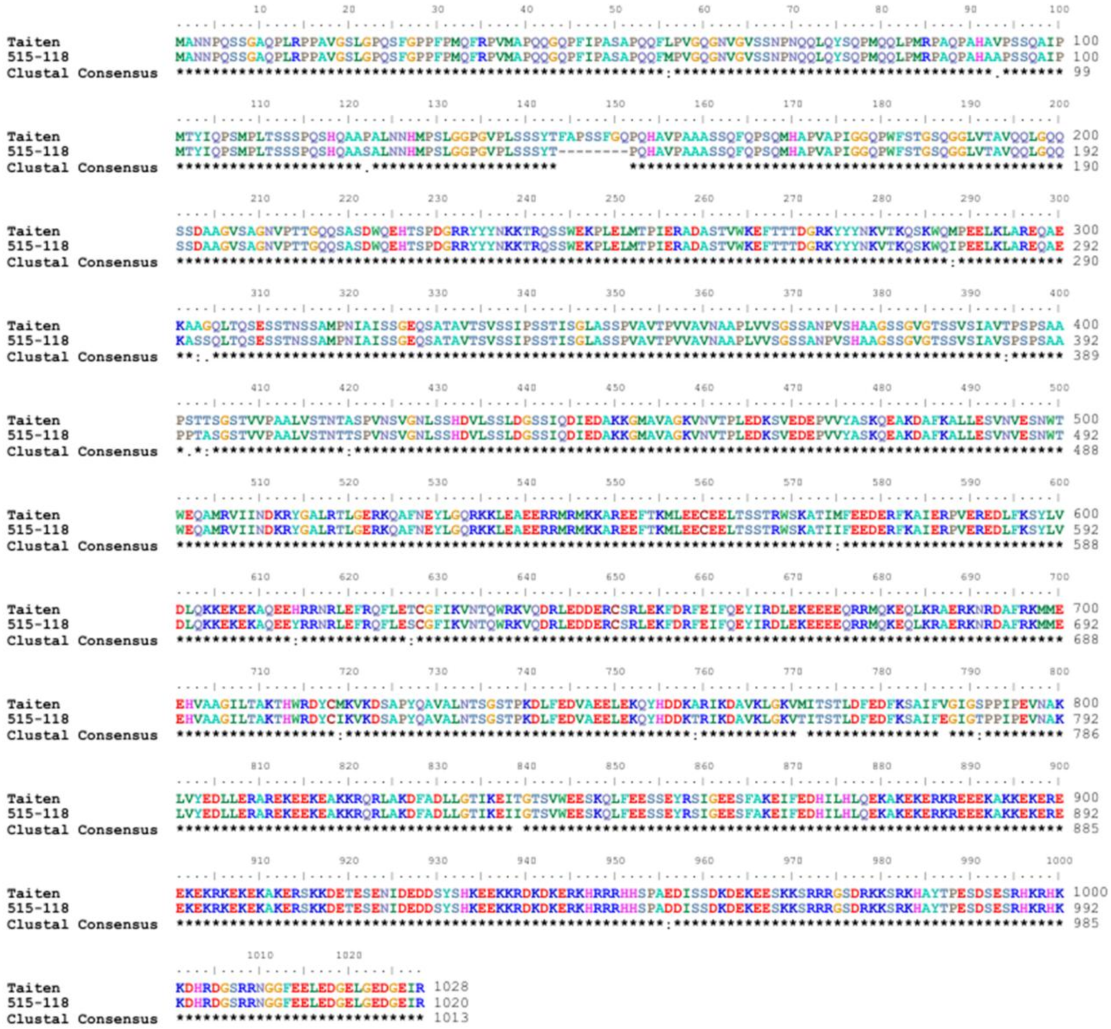

*PSK2*

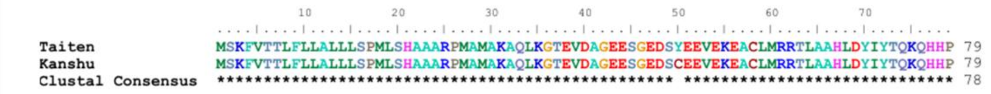

Unknown gene

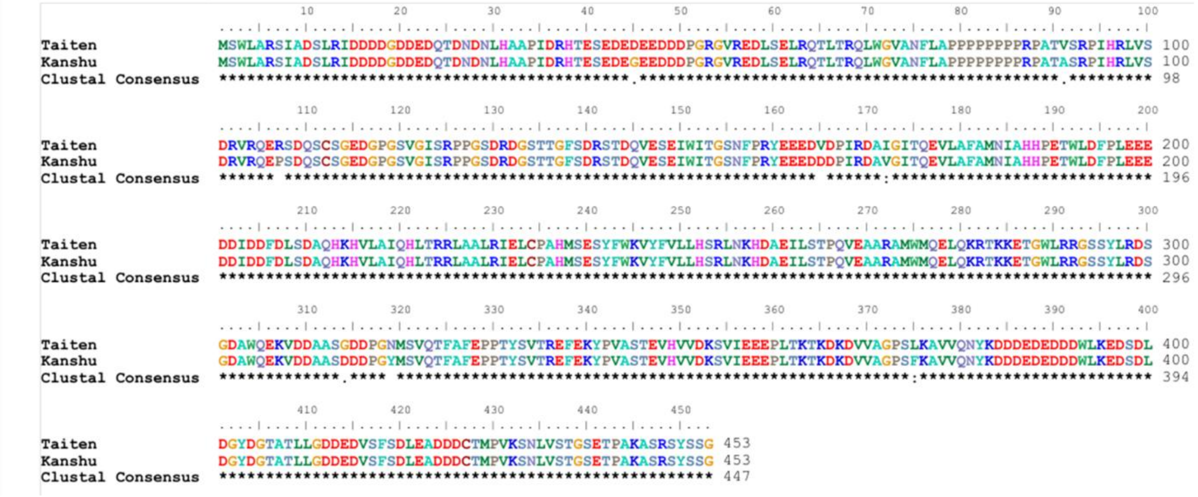

**Supplementary Fig. 10 Amio acid alignments of genes other than *DkTAN* that contain *AST*-linked polymorphisms**

Alignments were performed by ClustalW. Amino acid sequences were retrieved from Iso-Seq of ‘Taiten’ (non-PCNA) and ‘Kanshu’ (PCNA) or 515-118 (PCNA from ‘Taiten’ x ‘Kanshu’). No PCNA-specific abnormal transcripts were detected for the unknown gene, *PSK2* or *CIB4*. In *PRP40A*, the translated sequence from 515-118 contained a skipping region, but this alternation was not associated with *AST*-linked polymorphisms located in the intron.

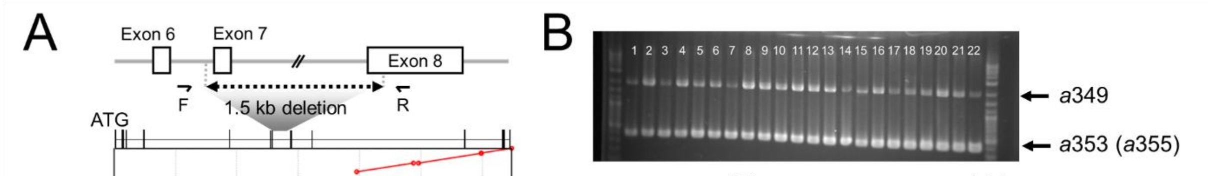


| | Cultivar | *ast* alleles | | | |
|---|---|---|---|---|---|
| 1 | Fuyu | 349 | 351 | 353 | 355 |
| 2 | Fukurogosho | 347 | 349 | 353 | 355 |
| 3 | Taishu | 349 | 351 | 353 | 355 |
| 4 | Fujiwaragosho | 349 | 353 | 355 | |
| 5 | Hanagohso | 349 | 351 | 353 | 355 |
| 6 | Shinshu | 347 | 349 | 353 | 355 |
| 7 | Soshu | 349 | 353 | 355 | |
| 8 | Jiro | 349 | 353 | 355 | |
| 9 | Tenjingosho | 349 | 353 | 355 | |
| 10 | Okugosho | 349 | 353 | 355 | |
| 11 | Mushhirodagosho | 349 | 353 | 355 | |
| 12 | Ogosho | 349 | 353 | 355 | |
| 13 | Tokudagosho | 349 | 353 | 355 | |
| 14 | Midai | 349 | 353 | 355 | |
| 15 | Suruga | 349 | 353 | 355 | |
| 16 | Mikado | 349 | 353 | 355 | |
| 17 | Taiga | 349 | 351 | 353 | 355 |
| 18 | Reigyoku | 349 | 351 | 353 | 355 |
| 19 | Kishu | 349 | 355 | | |
| 20 | Yoho | 349 | 351 | 353 | 355 |
| 21 | Taiho | 349 | 351 | 353 | 355 |
| 22 | Kanshu | 349 | 351 | 353 | 355 |

**Supplementary Fig. 11 Genomic structure and genotyping of the 1.5-kb deletion in *DkTAN***

(A) Comparison of haplotype sequences retrieved from the 'Taiten' assembly. Contigs were aligned to the *TAN* region in the *D. oleifera* genome. The gene model of *TAN* from *D. oleifera* annotation is shown above, and primer positions designed for panel B are also indicated (Forward: ACTCAACAAAGGCAAGTTTCTGTTATTAAG, Reverse: ATAATTAGAATTTTAATTCAGTTGGAATCAC). (B) Genotyping results for the 1.5-kb deletion in representative PCNA cultivars. Upper bands were amplified from the *a*349 haplotype, which lacks the deletion, whereas lower bands were amplified from the *a*353 or *a*355 haplotypes carrying the deletion. The table shows estimated *ast* haplotypes of each cultivar obtained from a previous report (*17*).

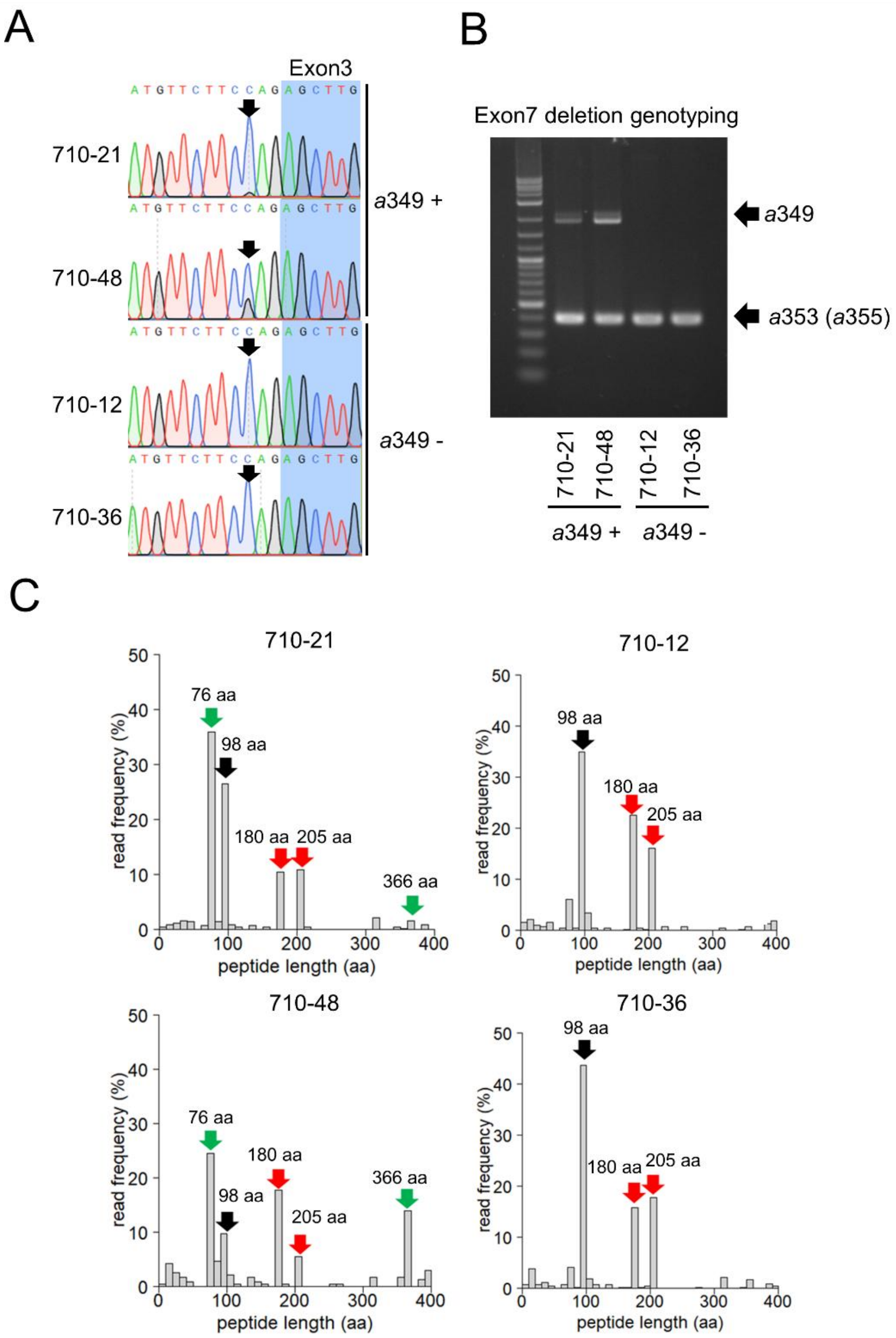


**Supplementary Fig. 12 Genotyping of the exon 3 SNP and exon 7 deletion, and cDNA amplicon sequencing of *DkTAN* in an *a*349-segregating population**

Four PCNA individuals (*aaaaaa*) from a segregating population derived from a cross between 'Midai' (PCNA) and 'Taiten' (non-PCNA) were analyzed. The presence or absence of the *a*349 allele was determined by fragment analysis as described by a previous report (*17*). (A) Sanger sequencing of PCR products spanning the exon 3 region. The arrow indicates the *a*349-specific SNP (C-to-G substitution) predicted to cause abnormal splicing. Individuals carrying the *a*349 allele (710-21 and 710-48) showed both C and G peaks, whereas individuals lacking *a*349 (710-12 and 710-36) showed only the C peak. (B) Genotyping of the exon 7 deletion. PCR was performed using the primer pair shown in Fig. S13. The *a*349-specific fragment was amplified only from individuals carrying the *a*349 allele. (C) Distribution of peptide lengths predicted from cDNA amplicon sequencing. Total RNA was extracted from immature fruits collected in early July, and library preparation and sequencing were performed as described in Materials and Methods. Black, green, and red arrows indicate the major peptide lengths encoded by *AST*, *a*349, and *a*353/*a*355, respectively. Transcripts derived from the *a*349 allele were detected only in individuals carrying *a*349.

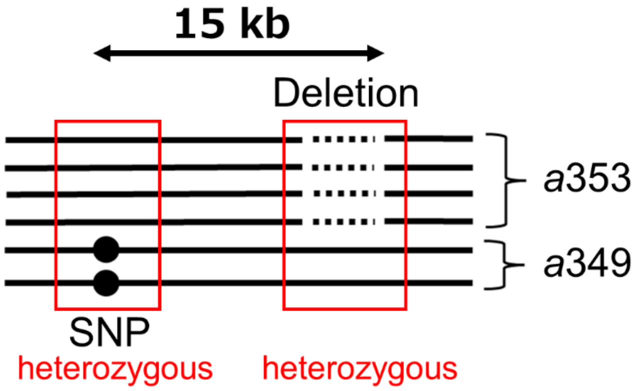


**Supplementary Fig. 13 Schematic illustration of the genomic positions of the two causal mutations in PCNA cultivars**

Because each mutation is present in only one of the two *ast* haplotype groups, genotyping based on either mutation alone results in an apparently heterozygous genotype, making PCNA cultivars indistinguishable from non-PCNA cultivars.

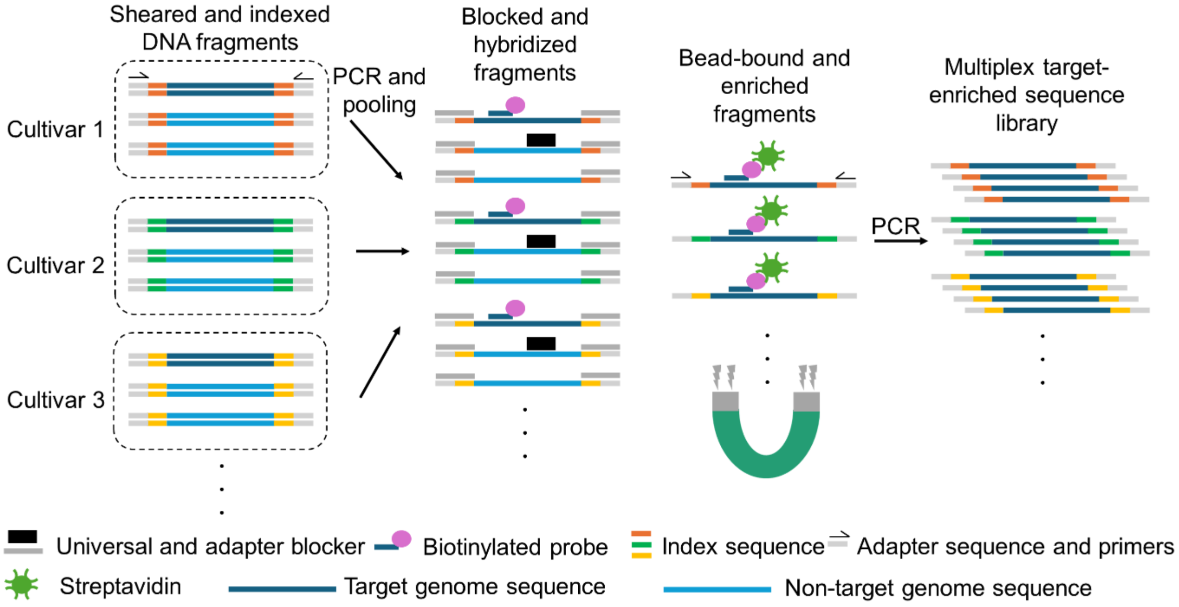


**Supplementary Fig. 14 Schematic overview of library construction for target-capture enrichment analysis**

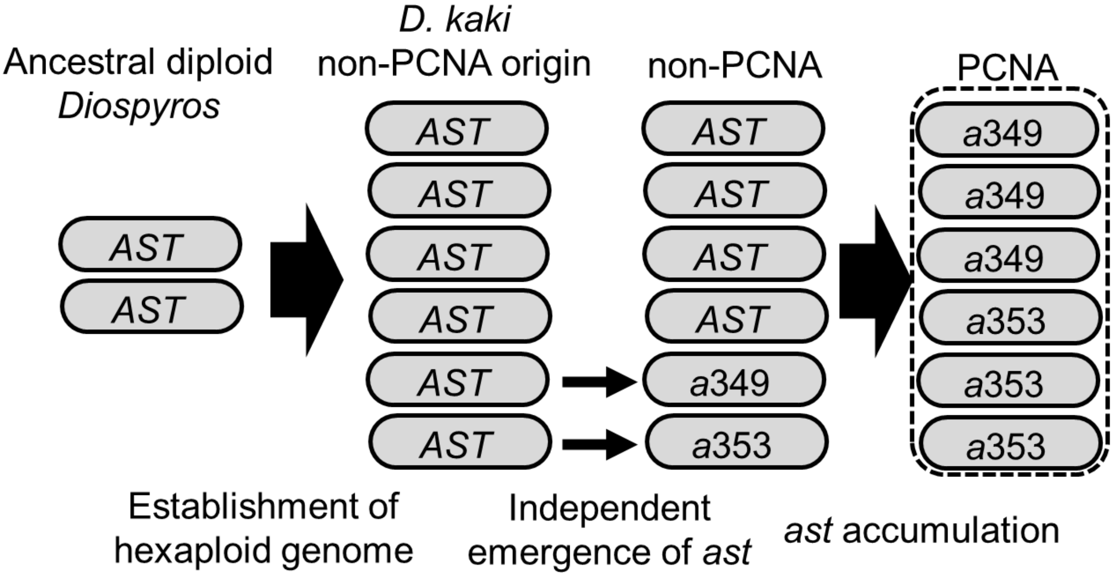


**Supplementary Fig. 15 Schematic model for the establishment of PCNA**

Haplotype network analysis suggested that the two *ast* groups, *a*349 and *a*355, were inferred to have arisen independently from ancestral *AST* haplotypes (or ancestral diploid alleles). PCNA is proposed to have been established through accumulation of six *ast* alleles belonging to the *a*349 and *a*355 groups.

### Supplementary Table 1 Summary of whole genome sequencing data

| Cultivar / ID | Class[*a] | Platform | Total sequencin g data (Gb) | Average read length (bp) | Sequencin g depth (x)[*b] | de novo genome asembly | Variant calling | Features | DRR Run Accession |
|---|---|---|---|---|---|---|---|---|---|
| Taiten | NP | Pacbio HiFi (7 cell) | 185.14 | 18,209 | 43.5 | ✓ | | | DRR1089512 - DRR1089518 |
| Taiten | NP | Illumina (pair end) | 65.13 | 130.4 | 15.3 | | ✓ | | DRR1089519 |
| Kurokuma | NP | Illumina (pair end) | 47.22 | 145.7 | 11.1 | ✓ | ✓ | Maternal parent of 'Taiten' | DRR1089520 |
| Tenryubo | NP | Illumina (pair end) | 102.48 | 148.7 | 24.1 | | ✓ | | DRR1089521 |
| Yoshidagosho | NP | Illumina (pair end) | 81.29 | 148.2 | 19.1 | | ✓ | Paternal parent of 652-line | DRR1089522 |
| Taishu | P | Illumina (pair end) | 64.93 | 148.7 | 15.3 | ✓ | ✓ | Paternal parent of 'Taiten' | DRR278876 |
| Fuyu | P | Illumina (pair end) | 78.5 | 143.2 | 18.5 | | ✓ | Maternal parent of 652-line | DRR278875 |
| Okugosho | P | Illumina (pair end) | 104.13 | 146.3 | 24.5 | | ✓ | | DRR1089523 |
| Kanshu | P | Illumina (pair end) | 61.42 | 129.8 | 14.4 | | ✓ | | DRR1089524 |
| Jiro | P | Illumina (pair end) | 102.52 | 146.4 | 24.1 | | ✓ | | DRR1089525 |
| Hanagosho | P | Illumina (pair end) | 105.57 | 148.1 | 24.8 | | ✓ | | DRR1089526 |
| Fujiwaragosho | P | Illumina (pair end) | 90.75 | 146.2 | 21.3 | | ✓ | | DRR1089527 |
| Mikado | P | Illumina (pair end) | 102.91 | 146.5 | 24.2 | | ✓ | | DRR1089528 |
| Midai | P | Illumina (pair end) | 100.3 | 146.7 | 23.6 | | ✓ | | DRR1089529 |
| 652-1 | NP | Illumina (pair end) | 22.1 | 146.9 | 5.2 | | ✓ | | DRR1089530 |
| 652-4 | NP | Illumina (pair end) | 27.31 | 147.7 | 6.4 | | ✓ | | DRR1089531 |
| 652-6 | NP | Illumina (pair end) | 28.49 | 147.5 | 6.7 | | ✓ | | DRR1089532 |
| 652-8 | NP | Illumina (pair end) | 30.76 | 148 | 7.2 | | ✓ | | DRR1089534 |
| 652-11 | NP | Illumina (pair end) | 24.2 | 146.1 | 5.7 | | ✓ | | DRR1089535 |
| 652-7 | P | Illumina (pair end) | 23.2 | 146.9 | 5.5 | | ✓ | | DRR1089533 |
| 652-16 | P | Illumina (pair end) | 32.53 | 147.7 | 7.7 | | ✓ | | DRR1089536 |
| 652-18 | P | Illumina (pair end) | 25.34 | 147.5 | 6 | | ✓ | | DRR1089537 |
| 652-31 | P | Illumina (pair end) | 27.46 | 148.2 | 6.5 | | ✓ | | DRR1089538 |
| 652-44 | P | Illumina (pair end) | 38.08 | 148.9 | 9 | | ✓ | | DRR1089539 |

*a : NP, non-PCNA: P, PCNA.

*b: Depth was calculated by dividing the total number of sequenced bases by the estimated size of the hexaploid *D. kaki* genome. The genome size was estimated based on the previously reported monoploid reference genome (*46*).

**Supplementary Table 2 Partition accuracy of trio-binning using parent-specific k-mers of different lengths**

| | Parent specific k-mer length | | | | | | | |
|---|---|---|---|---|---|---|---|---|
| | 17 | 19 | 21 | 23 | 25 | 27 | 31 | 37 |
| Reads accurately assigned for hapA (=a) | 90 | 91 | 92 | 93 | 93 | 96 | 96 | 55 |
| Reads accurately assigned for hapB (=b) | 41 | 64 | 73 | 80 | 99 | 74 | 90 | 0 |
| Partition accuracy (=a+b/254) | 0.5157 | 0.6102 | 0.6496 | 0.6811 | 0.7559 | 0.6693 | 0.7323 | 0.2165 |

A total of 254 Hi-Fi reads covering the *AST*/*ast* haplotype-discriminating region around Indel-3 were used in this analysis.

**Supplementary Table 3 Whole genome partitioning of HiFi reads using parent specific k-mers with k = 25 and k = 31**

| | Parent specific k-mer length | |
|---|---|---|
| | 25 | 31 |
| Reads assigned in hapA | 2,994,278 | 3,148,890 |
| Reads assigned in hapB | 3,390,302 | 4,289,332 |
| Reads assigned in hapU | 3,772,312 | 2,718,670 |

**Supplementary Table 4 Trio-binned assemblies using HiFi reads partitioned with k = 25 and k = 31**

| Hap. group | k | Total contig number | Length (bp) | | | |
|---|---|---|---|---|---|---|
| | | | min. | max. | Average | N50 (bp) |
| Contigs assembled from HapA + HapU group ('Kurokuma' haplotypes) | 25 | 4,391 | 445 | 39,075,432 | 548,730.90 | 6,885,818 |
| | 31 | 5,559 | 445 | 38,157,183 | 430,960.30 | 4,577,784 |
| Contigs assembled from HapB + HapU group ('Taishu' haplotypes) | 25 | 4,857 | 17,381 | 10,957,368 | 300,339.50 | 1,514,620 |
| | 31 | 6,108 | 445 | 26,400,783 | 417,855.50 | 5,728,044 |

### Supplementary Table 5 Gene annotation in the delimited *AST* region

| Position in the delimited *AST* region | | Annotation | | symbol |
|---|---|---|---|---|
| *A*224 haplotype | *A*240 haplotype | Gene ID in *D.oleifera* genome | Araport 11 blastn best-hit | |
| 559-36635 | -79870 | evm.model.fragScaff_scaffold_78.281 | endoplasmic reticulum-type calcium-transporting ATPase 3 (AT1G10130) | ECA3 |
| 140761-149688 | 133428-140070 | evm.model.fragScaff_scaffold_78.275 | no full name available (AT1G27480) | - |
| 168843-179115 | 141981-152115 | evm.model.fragScaff_scaffold_78.274 | no full name available (AT1G10720) | - |
| 184830-187351 | 157888-160783 | evm.model.fragScaff_scaffold_78.272 | phloem protein 2-A10 (AT1G10155) | PP2A10 |
| 223919-224433 | 200589-201103 | evm.model.fragScaff_scaffold_78.270 | phytosulfokine 2 precursor (AT2G22860) | PSK2 |
| 315216-348811 | 268374-301622 | evm.model.fragScaff_scaffold_78.264 | CW7 (AT1G59520) | CW7 |
| 481120-557265 | 421991-496003 | evm.model.fragScaff_scaffold_78.258_ evm.model.fragScaff_scaffold_78.259_ evm.model.fragScaff_scaffold_78.257 | pre-mRNA-processing protein 40A (AT1G44910) | PRP40A |
| 576187-601743 | 529433-551088 | evm.model.fragScaff_scaffold_78.256 | RNA-binding protein 1 (AT1G58470) | RBP1 |
| 624879-629210 | 573178-577050 | evm.model.fragScaff_scaffold_78.255 | CRY2-interacting bHLH4 (AT1G10120) | CIB4 |
| 687672-691218 | 638190-641665 | evm.model.fragScaff_scaffold_78.252 | DROUGHT HYPERSENSITIVE 2, SQUALENE EPOXIDASE 1 (AT1G58440) | SQE1 |
| 799669-827486 | 771225-791484 | evm.model.fragScaff_scaffold_78.248 | ZCW7 (AT1G59600) | ZCW7 |
| 849702-854015 | 817608-821483 | evm.model.fragScaff_scaffold_78.245 | no pollen germination related 1 (AT1G27460) | NPGR1 |
| 868731-870155 | 866816-868207 | evm.model.fragScaff_scaffold_78.241 | no full name available (AT5G43190) | - |
| 891401-892758 | 880729-881760 | evm.model.fragScaff_scaffold_78.239 | DWARF 14 (AT3G03990) | D14 |
| 970605-1006888 | 937275-964238 | evm.model.fragScaff_scaffold_78.230 | PLASTID ENVELOPE ATPASE 1, autoinhibited Ca2+-ATPase 1 (AT1G27770) | ACA1 |

The positions of genes on the *AST* haplotypes were determined using the haplotype-resolved sequences constructed in Fig. 1A. Gene annotations were assigned by homology searches of Iso-Seq-derived transcript sequences against the *D. oleifera* genome annotation (*19*) and the Araport11 database (*68*). The best-hit gene names or IDs are shown.

**Supplementary Table 6 Cultivars and *Diospyros* relative species used in full-length cDNA amplicon-seq of *TAN***

| PCNA | | | non-PCNA | | | Other *Diospyros* |
|---|---|---|---|---|---|---|
| Fujiwaragosho | Aizumishirazu | Ichiryo | Nagara | Shinpei | Yoshidagosho | *D. oleifera* |
| Fuyu | Akazu | Inayama | Nanyosuishi | Shirotodamashi | Yoshino | *D. lotus* Shinanoissaigaki |
| Hanagosho | Amahyakume | Iwaseto | Niikura | Shogatsu | Yotsumizo | *D. lotus* Chichibuissaigaki |
| Hazegosho | Amayotsumizo | Izaemon | Nijugaki | Shojo | Yotsuyasaijo | *D. lotus* Budogaki |
| Izu | Aoso | Jisya | Nikoronashiba | Shozaemon | Yozaburo | *D. virginiana* |
| Izushiogosho | Atago | Johonshi | Nishimurawase | Suimyo | Zenjimaru | *D. rhombifolia* |
| Jiro | Atagobo | Kaibuchi | Obishi | Suishi | | |
| Kanshu | Benigosho | Kakiyamagaki | Ohachiya | Syakokushi | | |
| Kashiwaragosho | Beniwase | Karasumi | Okujisya | Taigetsu | | |
| Kishu | Bongaki | Keizanbanshi | Okumyotan | Taiten | | |
| Matsuzaki | Butsushigano | Kikuhira | Onaga | Taiwankoshi | | |
| Midai | Dejimaru | Kikumanju | Onihira | Taiwanshoshi | | |
| Mikado | Dennai | Koedagaki | Oshoro | Takeda | | |
| Mizugosho | Dojohachiya | Kojimawase | Oyotsumizo | Takura | | |
| Mushirodagosho | Egosho | Konashiba | Pashi | Tamopan | | |
| Ogosho | Emon | Koshuhyakume | Rasyotomoe | Tenryubo | | |
| Okugosho | Gohanshi | Kurogaki | Rokuzaemon | Tohachi | | |
| Shinshu | Hacchiuri | Kyara | Saburoza | Toyoka | | |
| Soshu | Hagakushi | Luotian-tianshi | Saijo | Tyagone | | |
| Suruga | Hiratanenashi | Meotogaki | Saisyo | Wakamiya | | |
| Taishu | Hokikoshi | Mie | Sakata | Wasamatsuo | | |
| Tenjingosho | Hokisuishi | Mikatanigosho | Sangokuichi | Wasemyotan | | |
| Tokudagoho | Hosyomaru | Mizushimagosho | Sanja | Yamagaki | | |
| Yamatogosho | Ibogaki | Monpei | Seihakuji | Yashima | | |
| Youhou | Ichidagaki | Muraya | Shibumyotan | Yokono | | |

### Supplementary Table 7 Long read sequence and target enrichment result

| Cultivar | Class[*a] | Average HiFi read length (bp) | Coverage for target region | Enrich ratio for 'Taishu' *DkTAN* genomic region(%) | DRR accession number |
|---|---|---|---|---|---|
| Kakiyamagaki | NP | 3,794 | 14,968 | 63.201 | DRR1089714 |
| Kurokuma | NP | 3,651 | 20,510 | 58.031 | DRR1089717 |
| Yoshidagosho | NP | 4,090 | 11,256 | 65.678 | DRR1089733 |
| Taiten | NP | 4,041 | 19,543 | 60.036 | DRR1089729 |
| Fujiwaragosho | P | 3,946 | 13,758 | 70.164 | DRR1089709 |
| Fuyu | P | 3,850 | 16,119 | 69.636 | DRR1089710 |
| Hanagosho | P | 3,721 | 13,098 | 73.163 | DRR1089711 |
| Hazegosho | P | 3,870 | 22,355 | 61.204 | DRR1089712 |
| Jiro | P | 3,851 | 19,507 | 65.52 | DRR1089713 |
| Kanshu | P | 3,931 | 13,739 | 66.574 | DRR1089715 |
| Kishu | P | 4,196 | 15,320 | 66.188 | DRR1089716 |
| Midai | P | 3,865 | 17,251 | 68.972 | DRR1089718 |
| Mikado | P | 3,906 | 17,742 | 63.492 | DRR1089719 |
| Mushirodagosho | P | 3,733 | 15,242 | 69.925 | DRR1089720 |
| Ogosho | P | 3,972 | 15,465 | 69.228 | DRR1089721 |
| Okugosho | P | 4,050 | 11,200 | 75.048 | DRR1089722 |
| Reigyoku | P | 4,267 | 10,574 | 69.885 | DRR1089723 |
| Shinshu | P | 3,730 | 20,536 | 65.636 | DRR1089724 |
| Soshu | P | 3,822 | 16,944 | 69.477 | DRR1089725 |
| Suruga | P | 3,832 | 11,385 | 75.354 | DRR1089726 |
| Taiga | P | 4,021 | 18,027 | 64.057 | DRR1089727 |
| Taishu | P | 4,251 | 11,842 | 67.952 | DRR1089728 |
| Tenjingosho | P | 3,751 | 7,882 | 73.239 | DRR1089730 |
| Tokudagosho | P | 3,965 | 12,635 | 73.047 | DRR1089731 |
| Yoho | P | 4,328 | 14,110 | 66.71 | DRR1089732 |

*a : NP, non-PCNA: P, PCNA.

### Supplementary Table 8 Primer sequences

| Primer | Sequence (5'->3') | Features |
|---|---|---|
| **Amplicon sequencing** | | |
| AmpSeq_universal-F | CGAACATGTAGCTGACTCAGGTCACATGCTTGGTGCTGG AGATGAGACCCCTTCTAG | Adapter sequence for 2nd PCR is underlined |
| AmpSeq_universal-R | CACTTGTGCAAGCATCACATCGTAGCTACAAGCTAACTG GAGGACTTCCCTTAAATAAAAGATCATGAG | |
| Index-F | XXXXXCGAACATGTAGCTGACTCAGGTCAC | XXXXX indicates random index |
| Index-R | XXXXXCACTTGTGCAAGCATCACATCGTAG | |
